# Evolutionary history of the Brachyury gene in Hydrozoa: duplications, divergence and neofunctionalization

**DOI:** 10.1101/2023.01.09.523299

**Authors:** Alexandra A. Vetrova, Daria M. Kupaeva, Tatiana S. Lebedeva, Peter Walentek, Nikoloz Tsikolia, Stanislav V. Kremnyov

## Abstract

Brachyury, a member of T-box gene family, is widely known for its major role in mesoderm specification in bilaterians. It is also present in non-bilaterian metazoans, such as cnidarians, where it acts as a component of an axial patterning system. In this study, we present a phylogenetic analysis of Brachyury genes within phylum Cnidaria, investigate differential expression and address a functional framework of Brachyury paralogs in hydrozoan *Dynamena pumila*.

Our analysis indicates two duplication events of Brachyury in the cnidarian lineage: in the common ancestor of the Medusozoa clade and at the base of the class Hydrozoa. We designate result of the first step as Brachyury2 and of the second as Brachyury3.

Brachyury1 and 2 display a conservative expression pattern marking the oral pole of the body axis in *D. pumila*. On the contrary, Brachyury3 expression was detected in scattered presumably nerve cells of the *D. pumila* larva. Pharmacological modulations indicated that Brachyury3 is not under regulation of cWnt signalling in contrast to the other two Brachyury genes. Divergence in expression patterns and regulation suggest neofunctionalization of Brachyury3 in hydrozoans.

## Introduction

Brachyury (or T) is a founding member of T-box transcription factor family (Sebe-Pedrós, Ruiz-Trillo, 2016) first identified in a mutant mouse strain (Dobrovolskaia-Zavadskaia, 1927). Mice lacking one allele of Brachyury exhibit a short-tail phenotype (Smith, 1997), while the prenatal lethal loss of both alleles leads to severe deficiencies in mesoderm and axial structure formation (Gluecksohn-Schoenheimer, 1944; Gruneberg, 1958). Subsequent studies demonstrated that Brachyury is highly conserved and present not only in chordates, but in most metazoan animals ranging from ctenophores to sea urchins, as well as in ichthyosporeans, filastereans, and several early-branching fungi (Harada et al., 1995; Technau, 2001; Yamada et al., 2010; Sebe-Pedrós, Ruiz-Trillo, 2013).

Brachyury plays a crucial role in notochord formation in various chordates (reviewed in Satoh et al., 2012) and mesoderm specification in bilateria in general (reviewed in Bruce, Winklbauer, 2020), and its evolutionary primary function is possibly associated with germ layer demarcation and morphogenesis during gastrulation (Yasuoka et al., 2016; Lapebie et al., 2014). It is also an important component of the axial patterning gene regulatory network (Schwaiger et al., 2021).

Though functions of Brachyury were examined in a limited number of species, patterns of its expression during embryonic development are well studied. One of the Brachyury expression domains is conservatively detected at one pole of the body axis (e. g., oral pole in cnidarians, posterior pole in deuterostomes) (Kiecker, Niehrs, 2001; Peter, Davidson, 2011; Darras et al., 2018; Bagaeva et al., 2020), where the site of cell internalization is also located in the gastrulae of most animals. Within hydrozoans, Brachyury expression was demonstrated in the site of cell ingression during gastrulation in the embryos of *Clytia hemisphaerica* (Kraus et al., 2020). Brachyury is expressed around a blastopore in ctenophores (Yamada et al., 2010), anthozoans (Scholz & Technau, 2003; Yasuoka et al., 2016), echinoderms (Croce et al., 2001; Shoguchi et al., 2000), amphioxus (Yuan et al., 2020) and all vertebrates investigated so far (reviewed in Satoh et al., 2012), though this expression domain was lost in ascidians (Yasuo and Satoh, 1994). In annelids, mollusks, and insects Brachyury expression is also associated with the blastopore, though this expression domain dissolves to various degrees (reviewed in Bruce, Winklbauer, 2020).

A single copy of the Brachyury gene is present in genomes of most Metazoans. However, there are several exceptions. Within chordates, *Xenopus laevis* has four Brachyury genes (Latinkic et al., 1997; Hayata et al., 1999), where tbxt.L/tbxt.S (Xbra and Xbra2) and tbxt.2.L/tbxt.2.S (Xbra3) each are considered to be alloalleles, arising from the recent genome duplication (Latinkic et al., 1997; Hayata et al., 1999; Session et al., 2016). *X. tropicalis* contains two Brachyury genes, one of which is clustered with Xbra/Xbra2 (tbxt) and the other corresponds to Xbra3 gene of *X. laevis* (tbxt.2) (Martin and Kimelman, 2008). Teleost fish, such as medaka, zebrafish and three-spined stickleback, possess two Brachyury genes (Bra and Ntl) in their genomes (Halpern et al., 1993; Martin and Kimelman, 2008). Brachyury is present in two copies in the basal chordate amphioxus (Terazawa and Satoh, 1995, 1997; Yuan et al., 2020). According to phylogenetic analysis, duplication events occurred not in the chordate ancestor, but in all three chordate lineages independently (Inoue et al., 2017; Martin and Kimelman, 2008). Among non-chordate metazoans, the hydrozoans *Hydra* and *C. hemisphaerica* have at least two copies of the Brachyury gene (Bielen et al., 2007; Lapébie et al., 2014).

Thorough phylogenetic analysis is required to understand the evolution of Brachyury genes within cnidarians, in particular, whether gene duplication occurred in the common hydrozoan ancestor or if there were several independent lineage-specific events. To resolve this issue, we aimed to reconstruct the phylogenetic tree of Brachyury genes within phylum Cnidaria. Our data indicate a first gene duplication in the common ancestor of Meduzozoa. Strikingly, Brachyury has undergone one more duplication in the hydrozoan lineage, where we found three paralogs of Brachyury in most species. Next, analysis of gene expression patterns of Brachyury paralogs in the hydrozoan *Dynamena pumila* during normal development and in the colony demonstrated very different expression dynamics of DpBra3 from the expression of DpBra1 and DpBra2. Since it is known, that Brachyury is a direct target gene of the cWnt pathway (Vonica and Gumbiner, 2002; Bielen et al., 2007), we tested, if all three Brachyury paralogs are still under regulation of cWnt signaling. Data obtained from pharmacological modulations demonstrate that DpBra3 is differently regulated in comparison with DpBra1 and DpBra2. Taken together, our results suggest that the duplication of Brachyury genes in the hydrozoan lineage resulted in the diversification of the most recent copy.

## Results

### Diversity and phylogeny of cnidarian brachyury genes

To address the evolution of the Brachyury gene family within cnidarians, first we conducted TBLASTX search of the previously published transcriptome of *D. pumila* (Kupaeva et al., 2020) with the published *C. hemisphaerica* Brachyury gene sequences as an initial query. We recovered three sequences of Brachyury-like genes from the *D. pumila* transcriptome and used them as queries for TBLASTX searches against 10 more medusozoan transcriptomes (see Methods). Together with four already known anthozoan sequences, a total of 33 Brachyury sequences were identified from 16 Cnidaria species.

A maximum likelihood tree was generated using translated amino acid sequences with the best-fit JTT++R5 model (Figure 1). For this analysis, a total of 41 Brachyury sequences were used representing all major metazoan groups except Porifera. Since the T-box transcription factor family includes classes of Tbx genes besides Brachyury (Sebé-Pedrós et al., 2013), sequences of metazoan Tbx genes were used as an out-group to root the tree.

**Figure 1.**
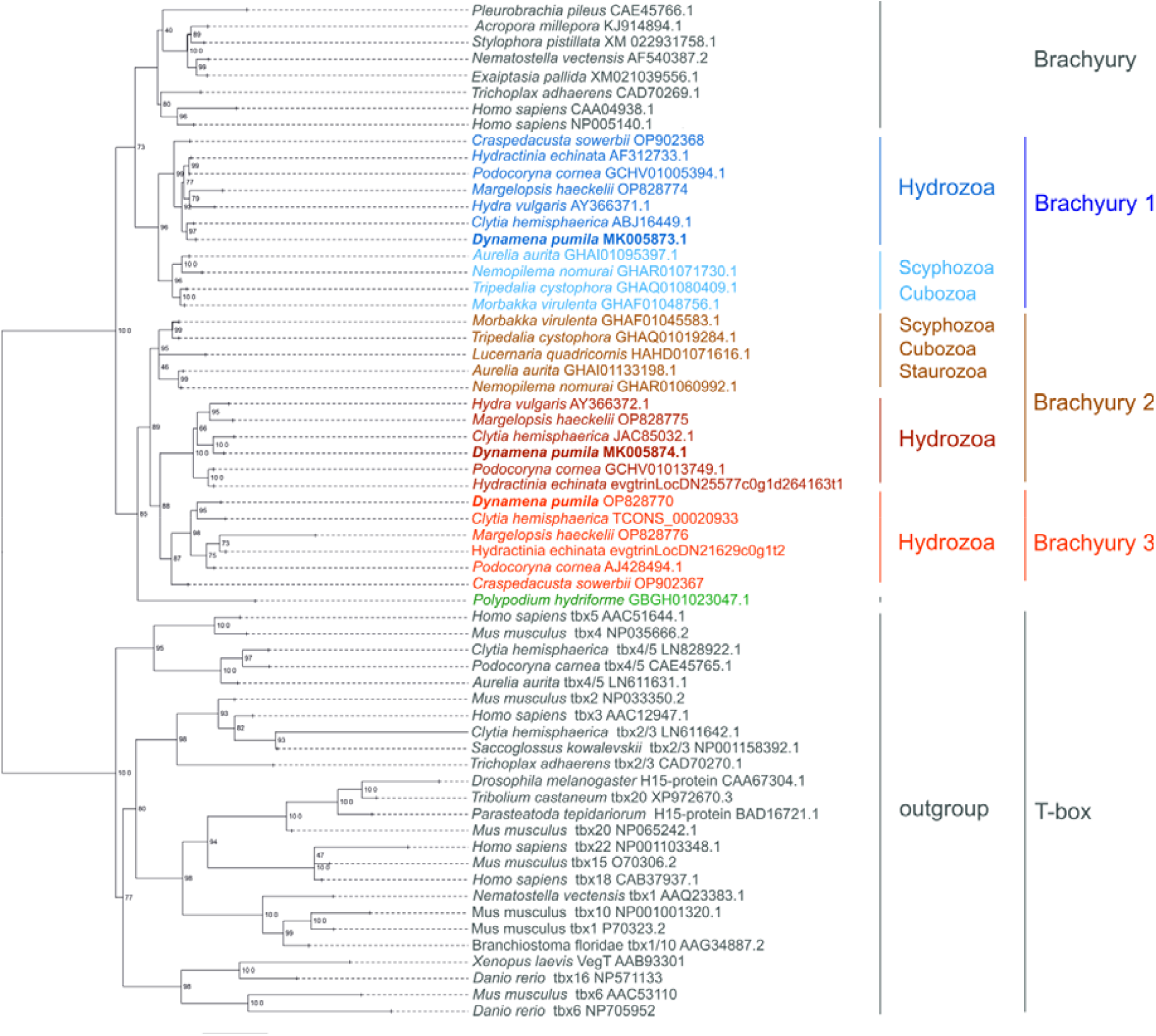
ML phylogenetic tree of Brachyury family members, rooted with TBX genes. Numbers at nodes are bootstrap values, shown as percentages. Scale indicates expected amino acid substitution per site. *D. pumila* genes are in bold.

All analyzed anthozoan species possess a single Brachyury gene (Figure 1). Anthozoan Brachyury sequences cluster with Brachyury genes of other not-cnidarian metazoans and form the sister group to the clade of medusozoan Brachyury1 genes. Besides Brachyury1, our transcriptomic survey revealed more Brachyury genes within Medusozoa (Figure 1). They cluster together as a sister group to Brachyury/Brachyury1 clade with a high nodal support (100% bootstrap value). Medusozoan-only Brachyury genes divide into cubozoan, scyphozoan and hydrozoan lineages. Whereas cubozoan and scyphozoan Brachyury2 genes cluster together, hydrozoan-only Brachyury genes form two sister groups, Brachyury2 and Brachyury3, with a well-supported bootstrap value (85%) (Figure 1). Interestingly, a previously studied Brachyury transcript (AJ428494.1) of *Podocoryne carnea* (Spring et al., 2002) is orthologous to Brachyury3 according to our analysis. Thus, all analyzed hydrozoan species have three Brachyury genes, with the exception of *Craspedacusta sowerbii* and *Hydra vulgaris*, which lack of Brachyury2 and Brachyury3, respectively.

### Comparison of sequence conservation of hydrozoan Brachyury proteins

Multiple sequence alignment of the deduced full-length amino acid sequences of *D. pumila* Brachyury proteins revealed that their T-boxes show about 70-77% identity. By contrast, full-length sequences have an overall lower amino acid identity, thus, the remaining regions are less conserved (Figure 2a, b).

**Figure 2.**
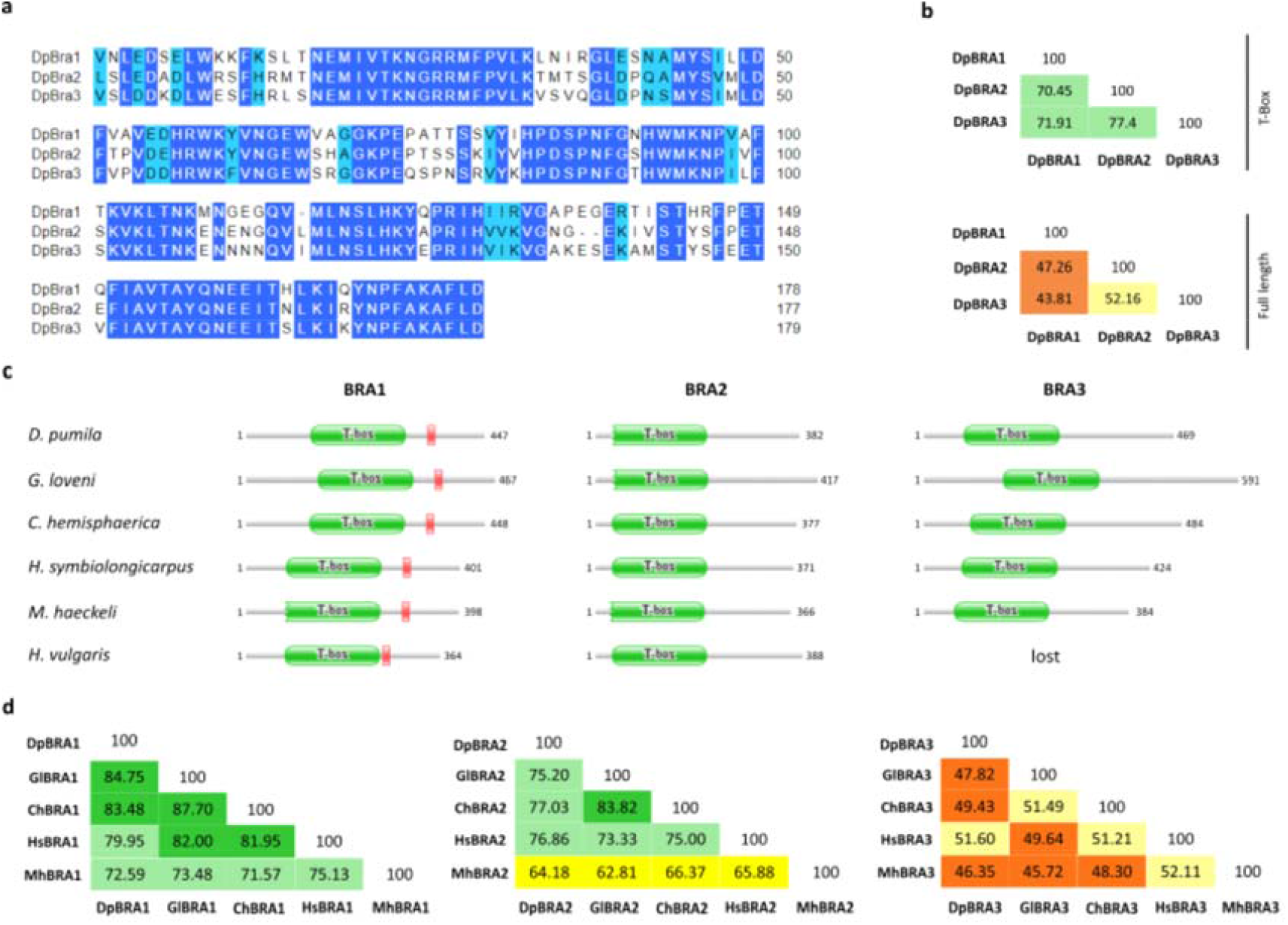
Comparison of sequence conservation among deduced hydrozoan Brachyury proteins. (a) Alignment of T-boxes of *D. pumila* Brachyury proteins. Amino acid identities are blue, light blue indicates that the residue is not identical but at least similar to the column consensus. (b) Percent identity matrix of T-boxes and full length Brachyury proteins in *D. pumila*. Colours here and below represent bands of percent identity: orange, 40-49%; light yellow, 50-59%; bright yellow, 60-69%; mint, 70-79%; green, 80-89%. (c) Domain architecture of predicted hydrozoan Brachyury proteins. A jagged edge indicates that a sequence match does not match the full length of the HMM that models a pfam entry. The number indicate the length of the protein in amino acids. (d) Percent identity matrix of full length hydrozoan Brachyury proteins.

Further, we performed a multiple sequence alignment of deduced hydrozoan Brachyury proteins, functional domain prediction by *hmmscan*, and a search of the R1 repressor domain with a sequence of R1 from *H. vulgaris* Bra1 as query (Bielen et al., 2007). Hydrozoan Bra1 proteins share higher identities with homologous genes in different species than Bra2 and Bra3 (Figure 2c, d). The inter-species Bra1 identities were also higher than identities between Brachyury paralogs within the same species (Figure 2b, d). Only Bra1 proteins have the repression R1 fragment in the C-terminal region (Figure 2c). Inter-species sequence comparison further revealed that of all analyzed hydrozoan Brachyury proteins, Bra2 proteins have the shortest sequences positioned N-terminally of the T-box (e.g., 20 amino acid for DpBra2), whereas Bra3 proteins are the most diverse in length and amino acid identity among hydrozoans (Figure 2c, d).

### Brachyury gene expression patterns during embryonic development and in shoots of the *D. pumila* colony

To determine whether the Brachyury paralogs are differently expressed during the development of *D. pumila*, we analyzed the spatiotemporal distribution of their transcripts by whole-mount *in situ* hybridization.

In situ hybridization revealed expression of DpBra1 in a unitary broad domain (Figure 3a) at the end of gastrulation, which does not overlap with any specific region within the gastrula stage embryo. Signal was visualized both in the ectoderm and the endoderm (Figure 3b, c). At the preplanula stage, expression signal was detected in discrete patches both in the ectoderm and the endoderm mostly at the oral end of the embryo (Figure 3d, e). In the early planula, we observed DpBra1 expression in the oral third of the larva (Figure 3f, g). In the mature planula, DpBra1 was expressed in the oral half of the larva (Figure 3h, i). In the ectoderm, we observed two domains of DpBra1 expression. In the tip, biased towards the oral pole expression signal was visualized in apical domains of ectodermal cells. Also, DpBra1 expression was visible in scattered ectodermal cells in the middle of the larva. In the endoderm, expression was present in only a few cells at the oral end (Figure 3j). Figure 3k represents expression patterns of DpBra1 during development.

**Figure 3.**
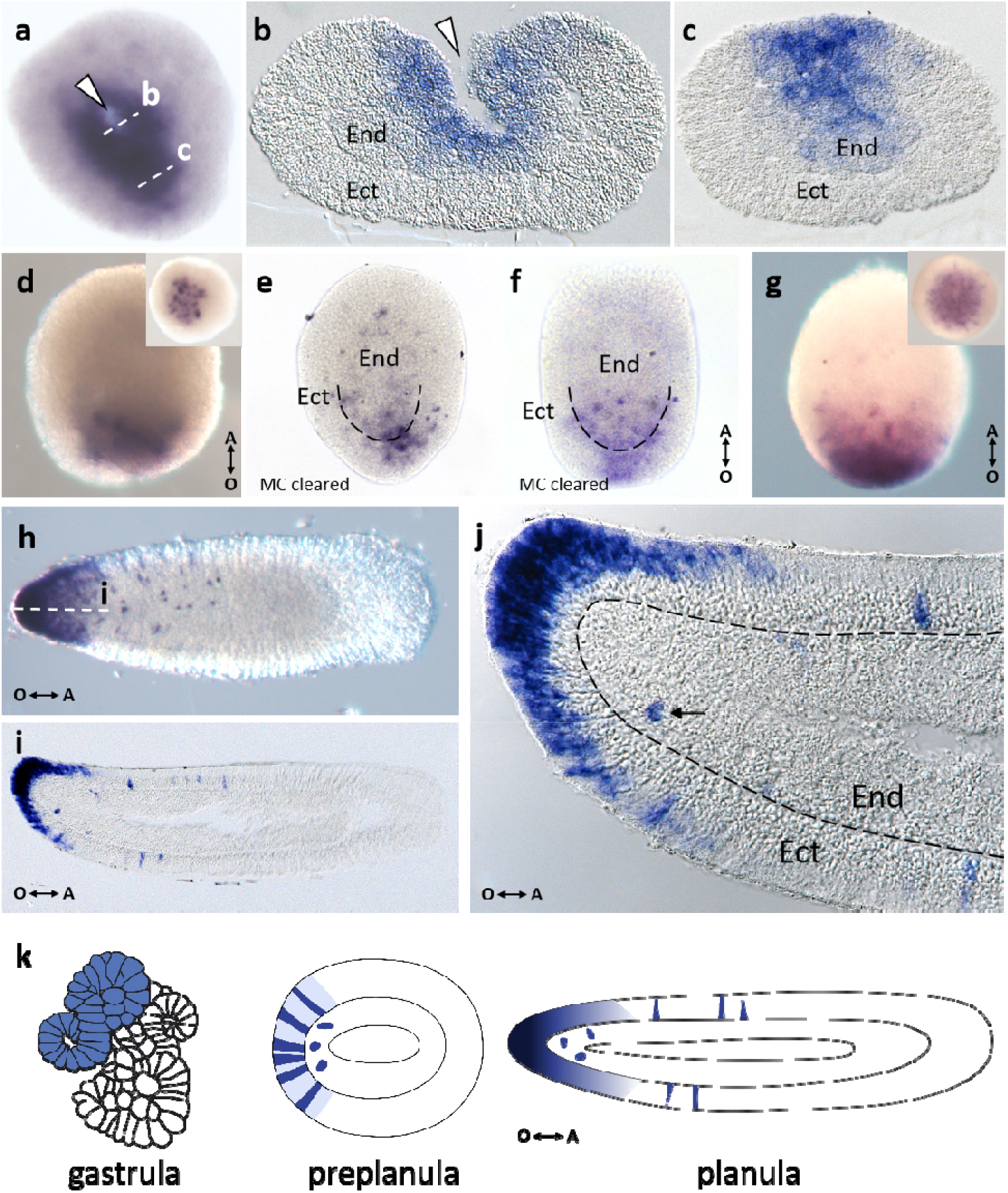
Spatial expression patterns of DpBra1 during embryonic development. (a) Expression is apparent around the epithelial torus in a broad domain at the end of gastrulation. White arrowhead points to the opening of the torus channel. (b, c) Transverse sections of the embryo through the levels indicated by the white dotted lines in a. Expression is present both in the ectoderm (Ect) and the endoderm (End). (d) Expressing cells are located at the oral pole of the preplanula. Double arrow shows the direction of the oral-aboral (O-A) body axis. (e) DpBra1 signal is prominent in the ectoderm and the endoderm of the preplanula cleared with Murray’s Clear (MC) solution. (f, g) Broad domain of expression biased towards the oral pole in the early planula. (h) In the mature planula, expression is observed in the oral end and in individual cells in the oral half of the body. (i) Longitudinal section of the planula through the level indicated by the white dotted line in h. Expression is localized in the oral ectoderm almost exclusively. (j) A blowup of the i. Black arrow indicates single endodermal cell with DpBra1 expression. Black dotted line marks the basal lamina.

DpBra2 expression was detected both in the ectoderm and the endoderm at the end of gastrulation and also forms a broad domain (Figure 4a, b). In the preplanula/early planula, DpBra2 expression was observed at the oral half of the embryo with more prominent signal at the oral pole (Figure 4c). Transcripts concentrated in the perinuclear cytoplasm of ectodermal cells (Figure 4d). In the mature planula, we observed DpBra2 expression in oral third of the larva with a bias towards the pole (Figure 4e, f). DpBra2 RNA was visualized in apical domains of ectodermal cells (Figure 4g). Expression was also present in single endodermal cells (Figure 4g) as in case of DpBra1 (Figure 3j). Figure 4h represents expression patterns of DpBra2 during development.

**Figure 4.**
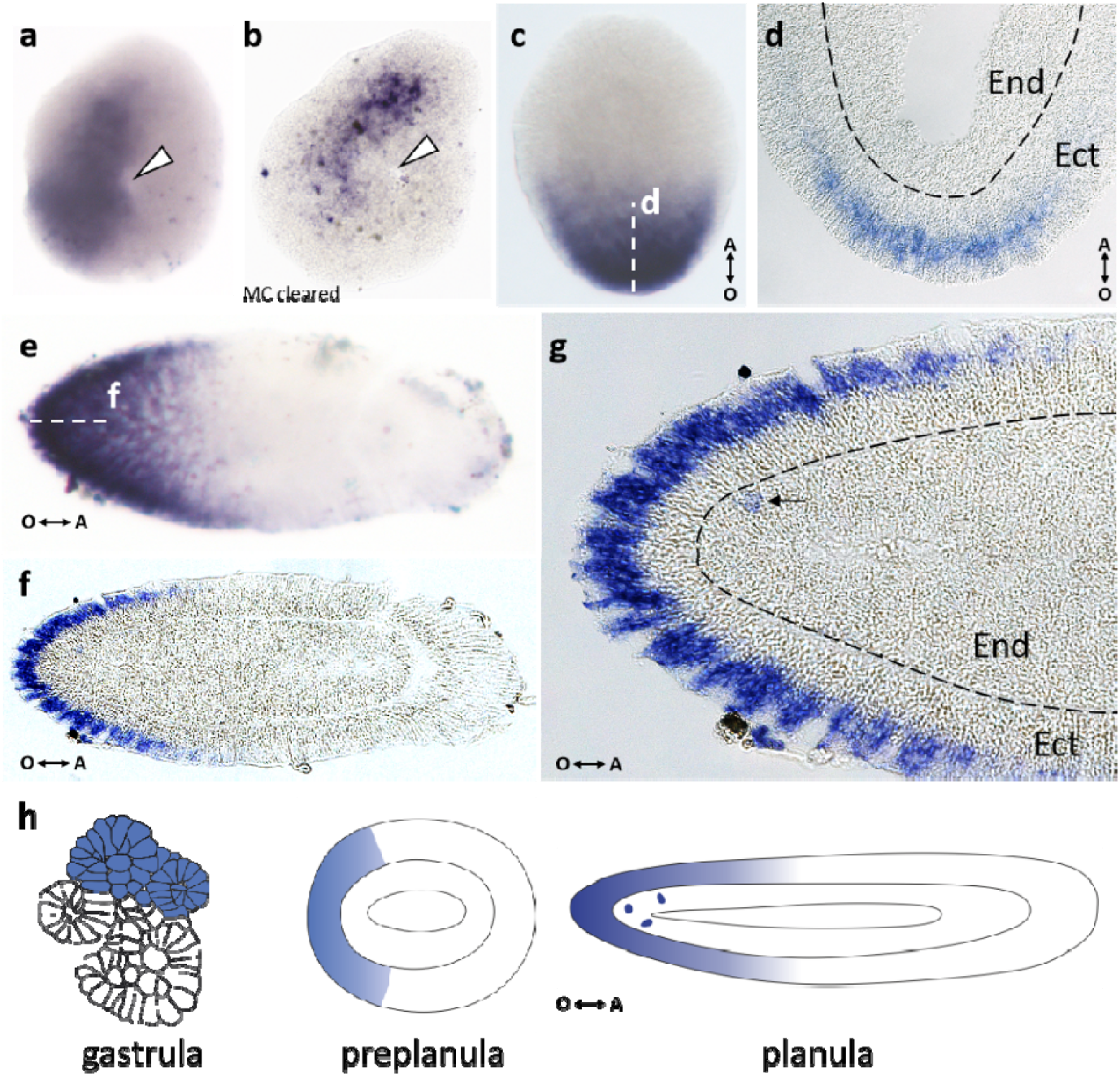
Spatial expression patterns of DpBra2 during embryonic development. (a) Broad expression domain is apparent near the last epithelial torus at the end of gastrulation. White arrowhead points to the opening of the torus channel. (b) Gastrula cleared with Murray’s Clear (MC) solution. Expresson signal is detected in inner cells of the embryo. (c) Biased toward the oral pole expression covers half of the preplanula/early planula. Double arrow shows the direction of the oral-aboral (O-A) body axis. (d) Longitudinal section of the embryo through the level indicated by the white dotted line in c. DpBra2 transcripts are visualized in the perinuclear cytoplasm of ectodermal (Ect) cells. End - endoderm. Black dotted line shows the basal lamina. (e) Oral expression is slightly biased to the pole in the mature planula. (f) Longitudinal section of the planula through the level indicated by the white dotted line in e. Expression is localized mostly in the oral ectoderm. (g) A blowup of the f. DpBra2 transcripts are visualized in apical domains of ectodermal cells. Black arrow indicates single endodermal cell with DpBra1 expression. B ack dotted line marks the basal lamina.

DpBra3 was expressed in a broad domain at the end of gastrulation (Figure 5a) in a pattern similar to those of DpBra1 and DpBra2. However, the expression pattern of DpBra3 differed drastically at later developmental stages. At the preplanula stage, DpBra3 signal was visible as a belt in the center of the oral-aboral axis (Figure 5b, c). As the development proceeded, the expression area expanded to cover the central part of the early planula (Figure 5d). Longitudinal sections (Figure 5e, f) revealed that transcripts are present mostly in basal domains of scattered ectodermal cells. As the planula elongated, expression continued in the middle part of the larva in discrete ectodermal cells (Figure 5g, h). Weak signal also appeared in the aboral endoderm (Figure 5g, h). Longitudinal sections of the mature planula clearly demonstrated that bottle-like (Figure 5i) and triangular (Figure 5j) bodies of DpBra3-expressing cells were located directly above the basal lamina or are between endoderm and ectoderm. The latter probably migrate towards the ectoderm from endoderm (Figure 5h). Figure 5k represents expression patterns of DpBra3 during development.

**Figure 5.**
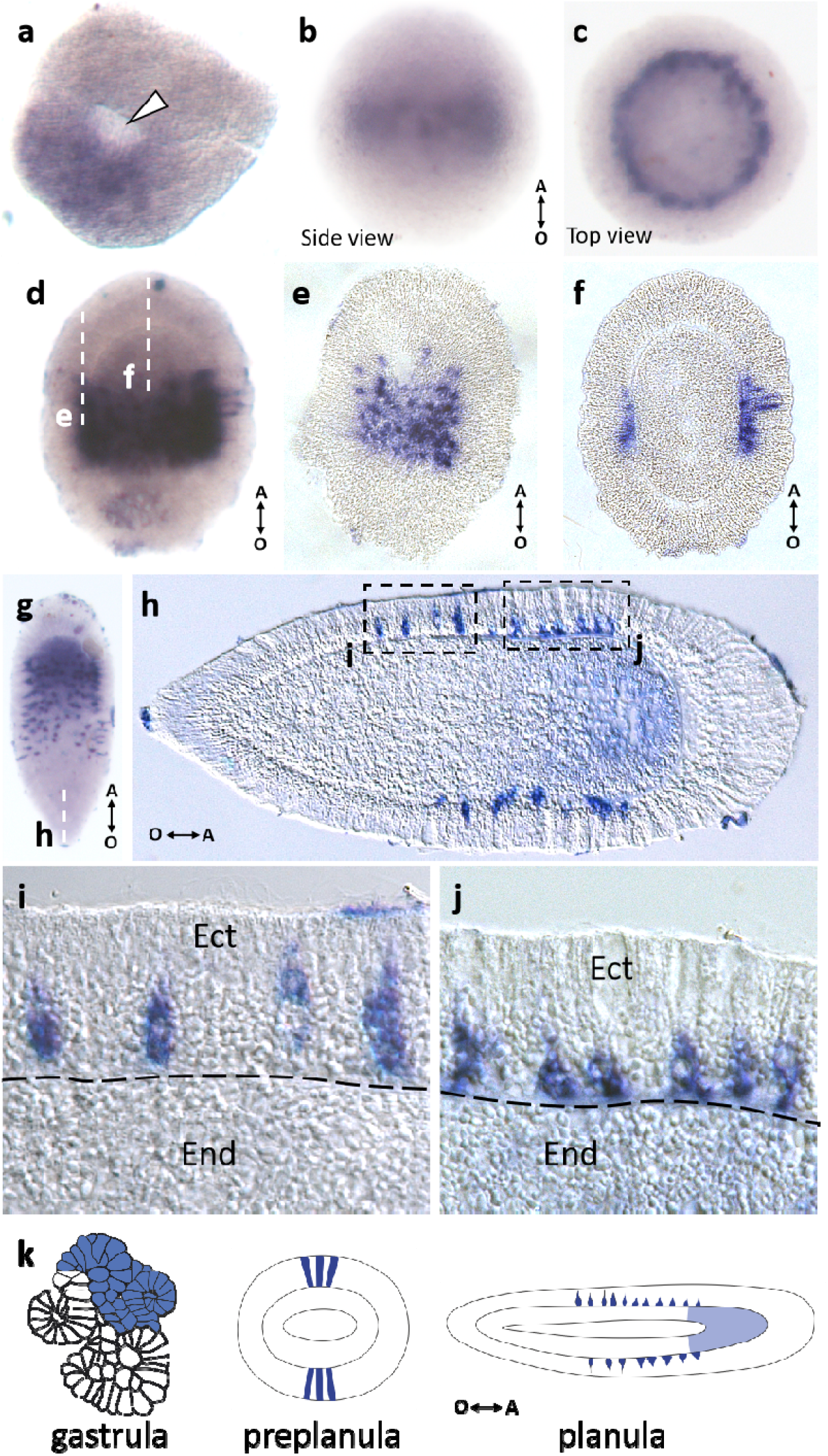
Spatial expression patterns of DpBra3 during embryonic development. (a) Broad expression domain is apparent near the last epithelial torus at the end of gastrulation. White arrowhead points to the opening of the torus channel. (b, c) Expression forms central belt showing in the ectoderm of the preplanula. Double arrow shows the direction of the oral-aboral (O-A) body axis. (d) Expressing cells are visible as a broad central belt in the early planula. (e, f) Longitudinal sections of the larva through the levels indicated by the white dotted lines in d. Expression is strictly ectodermal, staining is visualized mostly in basal cell domains. (g) Intense staining is visible in scattered cells in the central region of the mature larva. Weak staining is observed in the aboral endoderm. (h) Longitudinal section of the planula through the level indicated by the white dotted line in g. Intensely stained cells are located in the ectoderm. (i, j) Blowups of the h. Columnar and triangular bodies of expressing cells lie right above the basal lamina. Ect - endoderm, End - endoderm. Black dotted line marks the basal lamina.

Further, we examined expression patterns of the three Brachyury genes in the colony shoots of *D. pumila. D. pumila* forms monopodially growing colonies possessing biradial symmetry. Shoots of the colony are composed of repetitive modules. Each module consists of a fragment of the shoot in the center and two hydrants on the sides (Figure 6a). New modules are formed on the top due to the repeating morphogenetic cycle in the specific organ - the shoot growth tip (Figure 6b). Stage 1 represents the state when the morphogenetic cycle has not started yet (Figure 6b1). At stage 2, the growth starts with the apical surface of the tip curving up (Figure 6b2). At stage 3, the tip elongates and takes a hemispherical shape (Figure 6b3). At stage 4, the growth tip is dividing into the central and two lateral parts (Figure 6b4). Lateral primordia further differentiate into hydrants, while the central part will become the new shoot growth tip (Figure 6b1*).

**Figure 6.**
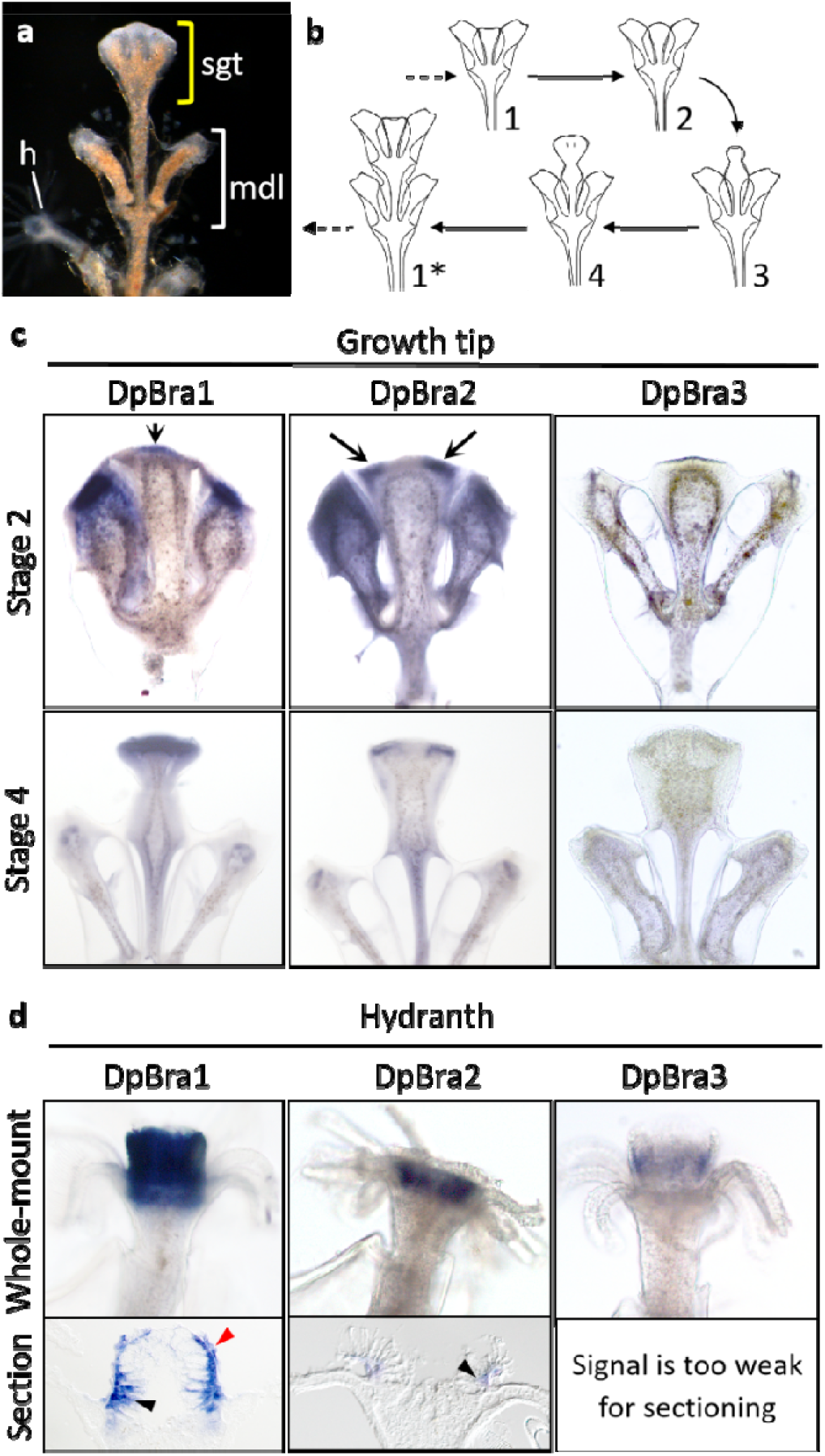
Spatial expresion patterns of brachyury genes in the colony of D. pumila. (a) The shoot of the D. pumila colony. Yellow bracket shows the shoot growth tip (sgt), white bracket - one module (mdl) of the shoot. h - hydranth. (b) The scheme of the morphogenetic cycle in the shoot growth tip of D. pumila. Numbers 1-4 indicate successive stages of morphogenesis. After the formation of the new internode, the cycle starts anew (*). (c) Spatial expresion patterns of brachyury genes in the shoot growth tips on stage 2 and 4 of the morphogenetic cycle. On the stage 2, DpBra1 expression is apparent in the central apical part of the shoot growth tip. DpBra2 expression is visible at the opposite sides of the shoot growth tip apex. On the stage 4, DpBra1 expression is uniform in the apex. DpBra2 expression remains at the opposite sides of the tip. Arrows point to the areas of expression. Expression of DpBra3 was not detected. (d) Spatial expresion patterns of brachyury genes in hydranths (whole-mount and londitudinal section through the center of the hydranth). Expression of brachyury genes is apparent in the hypostome of the hydranth. Black arrowheads point to expression in the endoderm. Red arrowhead points to expression in the ectoderm.

We analyzed expression patterns of three Brachyury genes in shoot growing tips on stages 2 and 4 of the morphogenetic cycle and in fully formed differentiated hydrants. DpBra1 and DpBra2 expression was detected in the apical ectoderm of the growth tip (Figure 6c). DpBra1 is expressed in the central part of the apex at stage 2 and uniformly at stage 4. DpBra2 expression was observed in two domains at opposite sides of the apex at stages 2 and 4. Thus, DpBra1 and DpBra2 expression domains do not overlap at stage 2, but are co-expressed at stage 4. DpBra3 expression was not detected in the shoot growth tip.

Expression of three Brachyury genes was observed in the hypostome of the hydranth (Figure 6d). A longitudinal section through the center of the hydranth revealed that DpBra1 was expressed both in the ecto- and the endoderm. DpBra2 signal was clearly visible in the endoderm of the hypostome, while the presence of a signal in the ectoderm is unclear. Unfortunately, DpBra3 signal was too weak for the fine examination, but seems to be expressed in the ectoderm (when viewed from the surface).

### Brachyury genes are differently regulated by the cWnt signaling in *D. pumila*

It was shown previously in *Hydra* and *C. hemisphaerica* that two hydrozoan Brachyury genes, Brachyury1 and Brachyury2, are regulated by cWnt signaling (Bielen et al., 2007; Momose and Houliston, 2007; Momose et al., 2008; Lapebie et al., 2014). However, it is unknown if Brachyury3 is still a cWnt-dependent gene after the duplication event. We assayed the dependence of three Brachyury genes on the cWnt pathway in *D. pumila*. We treated embryos at the gastrula stage with different concentrations of pharmacological agents to modulate the cWnt pathway, cultivated them until planula stage, and examined then expression patterns of three Brachyury genes in planula larvae of *D. pumila* (Figure 7). Azakenpaullone (Azk) activates cWnt signaling and iCRT14 inhibits it (Kunick et al., 2004; Stukenbrock et al., 2008; Gonsalves et al., 2011). It was shown in a previous study that hyper-activation of cWnt signaling results in the enlargement of larval oral domain, while its inhibition leads to reduction of oral domain in *D. pumila* (Vetrova et al., 2021).

**Figure 7.**
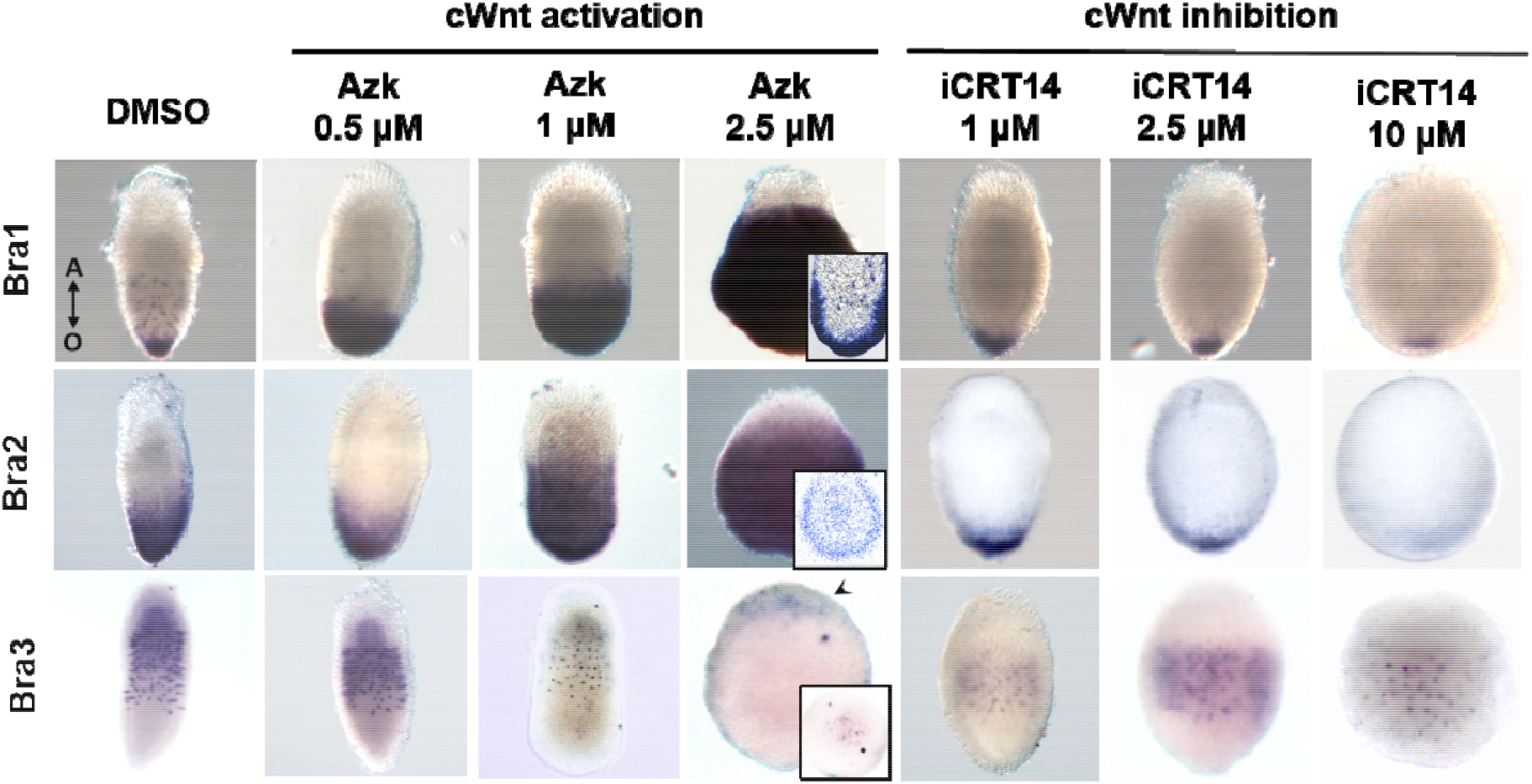
DpBra1 and DpBra2 expression depend on the activity of the cWnt signaling pathway. Pharmacological modulations of the cWnt pathway change the area of DpBra1 and DpBra2 expression in planula larvae. Number of endodermal signal-positive cells also increases (see details for longitudinal sections). Hyperactivation of the cWnt results in DpBra3 expression domain shifted aborally. cWnt inhibition does not effect DpBra3 expression notably. Arrowhead points to DpBra3-expressing cells on the oral pole of the larva (see detail). Double arrow shows the direction of the oral-aboral (O-A) body axis for all larvae.

DMSO-treated (control) larvae had normal morphology and expression patterns of three Brachyury genes (Figure 7). Treatments with the increasing concentrations of Azk resulted in the gradual expansion of DpBra1 and DpBra2 expression domains. After 2.5 μM Azk treatment, DpBra1 and DpBra2 expression signals were observed in the entire larva except the aboral-most region. The number of endodermal DpBra1- and Bra2-positive cells also increased. Vice versa, gradual inhibition of the cWnt signaling with iCRT14 led to the decrease of DpBra1 and DpBra2 expression domains in area (Figure 7).

Strikingly, overactivation of the cWnt signaling didn’t lead to the expansion of DpBra3 expression domain. The belt of DpBra3-expressing cells shifted in more and more aboral positions, vacating the central domain. In the result of 2.5 μM Azk treatment, several DpBra3-expressing cells were detected at the aboral pole of the larva (Figure 7: arrowhead). Inhibition of the cWnt signaling did not notably change the DpBra3 expression domain (Figure 7).

### *D. pumila* Brachyury genes differently regulate tissue differentiation in the animal cap assay

To uncover functional differences of three *D. pumila* Brachyury genes, we employed the *Xenopus laevis* animal cap assay system. Using this assay, we surveyed DpBra1, DpBra2, and DpBra3 for their ability to affect cell fates of naive *Xenopus* animal cap cells. It is known, that untreated animal caps differentiate into epidermal tissue (Green, 1999), but the injection with *Xenopus* Bra or *Hydra* Bra1 mRNA promotes mesoderm specification, and *Hydra* Bra2 mRNA shows neural-inducing activity (Smith et al.,1991; Bielen et al., 2007). We injected capped mRNAs encoding DpBra1, DpBra2, or DpBra3 into the animal region of two- to four-cell stage embryos (∼1 ng per embryo), dissected the animal caps at the blastula stage (stage 8), cultured them until control embryos reached late neurula stage (stage 18), and examined marker gene expression in these caps using conventional or quantitative RT-PCR (qRT-PCR). Uninjected animal caps were used as a control group.

DpBra1 significantly (P<0.0001) induced the expression of mesodermal marker gene actc1.L (muscle actin) (Figure 8a) as well the expression of another mesodermal marker gene, myod.S. Since myod.S expression was too low for reliable quantification in the control group using qRT-PCR, we used gel electrophoresis to show the induction (Figure 8c). DpBra1 did not affect neural differentiation (Figure 8b) while DpBra2 and DpBra3 did not show mesoderm- or neural-inducing activity (Figure 8).

**Figure 8.**
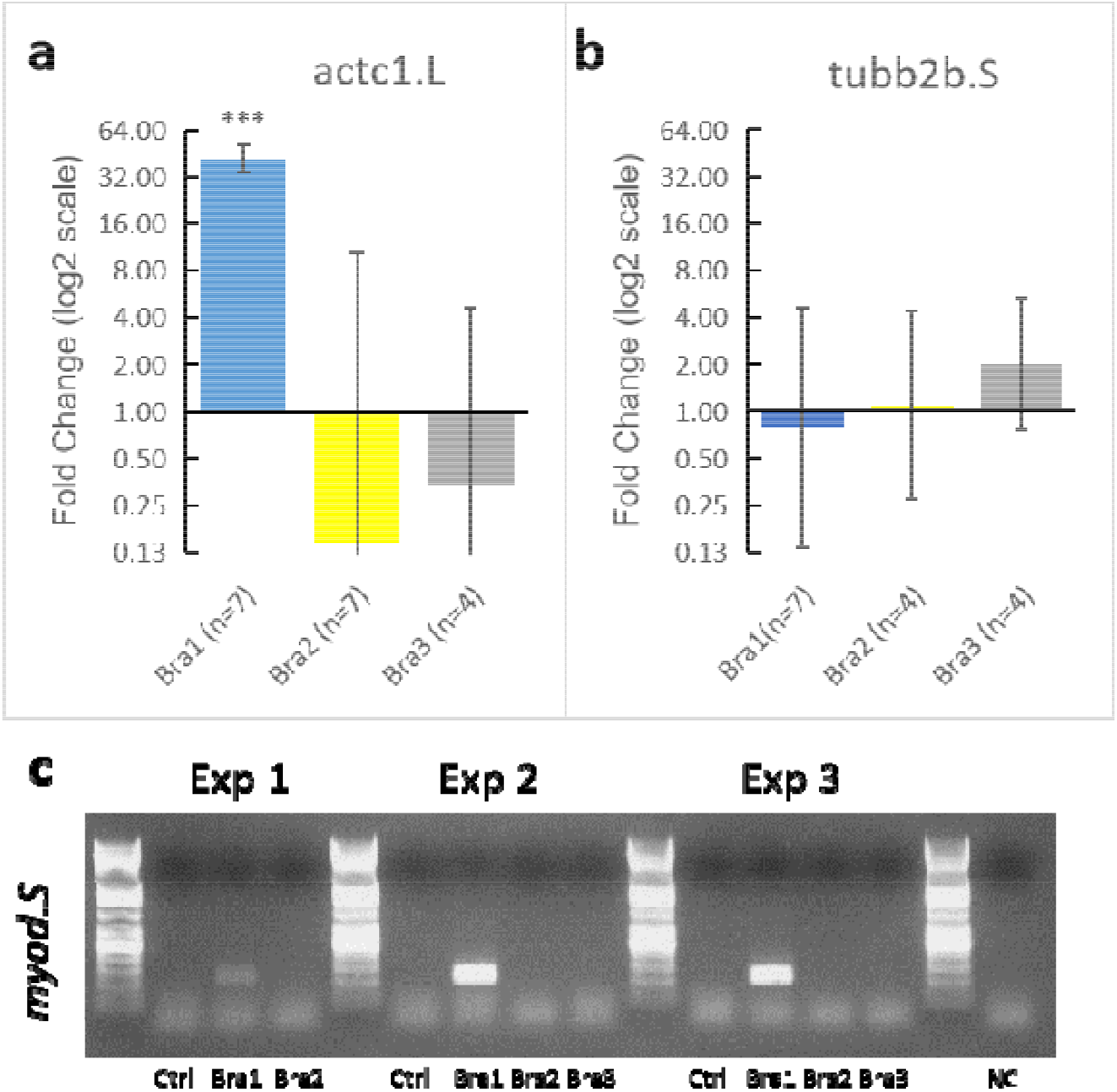
Molecular phenotype of *Xenopus* animal caps injected with *D. pumila* Brachyury genes. (a, b) RT-qPCR analysis on the induction of actc1.L, myod2.S, and tubb2b.S by *D. pumila* Brachyury mRNAs. Data are presented as normalized fold change expression (mean±s. d.) in experimental groups. n - n-value. The number of experimental groups is 3, 3, and 2 in (a); 3, 2, and 2 in (b). ***: p<0.001. (c) Gel electrophoresis for myod.S after injection by *D. pumila* Brachyury mRNAs. NC - negative control.

## Discussion

Gene duplications have long been recognized to facilitate the evolution of regulatory genes, driving expansion of families of signaling molecules and transcription factors (Gu, 2003; Wagner, 2008). Duplication event produces two copies (paralogs) of a gene both of which are orthologous to the “parent” gene. Subsequent evolution may lead to divergence: one copy retains high similarity to its orthologs while the second undergoes structural changes and may be designated as a daughter or child copy (Perry, Assis, 2016). In the present study, three paralogous genes of Brachyury transcription factor were found in hydrozoan lineage. The phylogenetic reconstruction suggests that one duplication event occurred first in the common ancestor of all meduzozoan species. The “daughter” copy of the first duplication (Brachyury2) experienced then an additional duplication in the common ancestor of hydrozoans (Figure 1).

There are three main scenarios after gene duplication. First outcome of gene duplication leads to loss of function of duplicated copy which becomes a pseudogene or is lost. Two other outcomes are subfunctionalization and neofunctionalization (Force et al., 1999; Lynch and Force, 2000). In case of subfunctionalization, two duplicated copies share the original function of the ancestral gene, and both are required to preserve the entire ancestral function (Force et al., 1999; Conant & Wolfe, 2008). Overlapping expression domains of two duplicates often reflect occurred subfunctionalization (Force et al., 1999). In case of neofunctionalization, one copy retains its ancestral functions, and the other one is free to gain a novel function, since it is relaxed from the selective pressure to perform its previous role (Ohno, 1970; Krakauer and Nowak, 1999; Conant & Wolfe, 2008). In this case, one duplicate acquires the expression domain different from the alternate copy and the ancestral gene (Gu et al., 2005; Assis, Bachtrog, 2013). The same duplicate can also display both features of sub- and neofunctionalization with regard to different functions (He, Zhang, 2005). The acquisition of novel functions by regulatory genes play a key role in diversification of developmental pathways and body plans in metazoans (Carroll, 2001; Larroux et al., 2008).

Evolution of Brachyury reveals signs of both sub- and neofunctionalization. In teleosts, two Brachyury genes are expressed in the marginal zone throughout gastrulation (Schulte-Merker et al., 1994; Martin and Kimelman, 2008) and subsequently in the notochord and the tailbud, revealing the common chordate pattern (Schulte-Merker et al., 1994; Martin and Kimelman, 2008). Only simultaneous loss of both Brachyury genes results in lack of somites and a notochord beyond the 12th somites (Martin and Kimelman, 2008), recapitulating mouse homozygous brachyury mutant phenotype (Gluecksohn-Schoenheimer, 1944) and indicating subfunctionalization followed duplication of Brachyury in teleosts. On the contrary, neofunctionalization seems to follow Brachyury duplication in *X. laevis* as XBra3 (tbxt2.L/tbxt2.S) is distinct from XBra and XBra2 (tbxt,L/tbxt,S) in function and spatio-temporal pattern of expression (Strong et al., 2000).

Within hydrozoans, outcomes of Brachyury duplication previously were studied in *Hydra* (Bielen et al., 2007). Though both Brachyury paralogs are expressed in the hypostome of *Hydra*, they evolved distinct coding sequences and diverged their functions. Authors posit, that Brachyury paralogs have undergone a mixture of sub- and neofunctionalisation in *Hydra* (Bielen et al., 2007). However, it is unknown, if similar outcomes have followed Brachyury duplication in other cnidarians.

As in *Hydra*, in other cnidarians, Brachyury genes are involved in molecular axial patterning (Schwaiger et al., 2021). In cnidarian species with polar gastrulation, that is, when it is linked to the axial patterning and occurs in the oral region of the embryo (Kraus, Markov, 2017), Brachyury expression accompanies gastrulation morphogenetic movements. Its expression was detected at the blastopore margin in anthozoan species (*Nematostella, Acropora*) (Scholz & Technau, 2003; Yasuoka et al., 2016). In hydrozoans *P. carnea* and *C. hemisphaerica*, gastrulation occurs not by invagination, but by unipolar cell ingression (reviewed in Kraus, Markov, 2017), yet Brachyury homologs are expressed in the respective region. In *C. hemisphaerica*, both studied Brachyury paralogs (ChBra1 and ChBra2) are expressed there with overlapping patterns (Kraus et al., 2020) and both are important for progression of gastrulation (Lapebie et al., 2014). Similarly, in *P. carnea* Brachyury was detected in the region of gastrulation on the oral pole (Spring et al., 2002) though the studied transcript is, according to our analysis, orthologous to Brachyury3 (Figure 1). In *D. pumila* during gastrulation, all three Brachyury paralogs are expressed in a broad domain (Figure 3a, 4a, 5a). In contrast to the cnidarian species with polar gastrulation (Yasuoka et al., 2016), it is unlikely, that Brachyury genes provide demarcation of ecto-endoderm boundary in *D. pumila*, since germ layers specification is not associated with axial polarity and oral region in particular during gastrulation in this species (Vetrova et al., 2021).

Predominantly oral expression of Brachyury genes continues throughout a cnidarian life cycle. In larvae of anthozoan species (*Nematostella, Acropora*) Brachyury is expressed in pharynx (Scholz & Technau, 2003; Yasuoka et al., 2016). In larvae of hydrozoans *C. hemisphaerica* and *D. pumila*, expression of Brachyury1 and Brachyury2 orthologs is detected in the oral ectoderm (Kraus et al., 2020) (Figure 3h-j, 4e-g). But unlike *C. hemisphaerica* where Brachyury orthologs are expressed in the oral ectoderm only, in *D. pumila* expression of DpBra1 and DpBra2 was also detected in a few scattered oral endodermal cells (Figure 3j, 4g). This expression pattern displays similarity to the expression of endodermal marker gene FoxA in *D. pumila* planula (Vetrova et al., 2021). Since Brachyury is conservatively co-expressed with FoxA in metazoans ranging from the anthozoan *Nematostella* (Servetnik et al., 2017) to a sea urchin (Tu et al., 2006) and *Drosophila melanogaster* (Kusch and Reuter, 1999), we infer that in *D. pumila*, these oral endodermal cells co-express Brachyury and FoxA as well.

Surprisingly, Brachyury3 orthologs are not associated with oral tissues in hydrozoan larvae and evolved to lineage-specific expression domains. Previously examined Brachyury ortholog of *P. carnea* clusters with Brachyury3 group according to our analysis (Figure 1), and its expression was detected not in the oral, but in the aboral ectoderm (Spring et al., 2002). In *D. pumila*, Brachyury3 expression also was detected in discrete triangular and bottle-like ectodermal cells (Figure 5g-j), morphologically similar to sensory cells of cnidarian nerve system (Grimmelikhuijzen & Westfall, 1995). This is of interest considering that Brachyury gene acts as neural repressor in the anthozoan *Nematostella* (Schwaiger et al., 2021). Moreover, DpBra3 seems not to be a cWnt-dependent gene in contrast to other cnidarian and bilaterian Brachyury genes studied in this regard (Arnold et al., 2000; Schwaiger et al., 2021), DpBra1 and DpBra2 included (Figure 6). Comparison of sequence conservation demonstrates that Brachury3 orthologs are the most diverse in length and amino acid identity among hydrozoan Brachyury gene family (Figure 2c, d). Differences in protein sequences, regulation, and expression domains suggest that the newly derived Brachyury3 diverged to different functions in hydrozoan species. It is common to the youngest paralog to acquire new functions since its ancestral function is already covered by older duplicates (Assis, Bachtrog, 2013).

In the hydranth of *D. pumila*, DpBra3 expression pattern does not drastically differ from patterns of DpBra1 and DpBra2 and is in line with previous studies (Spring et al., 2002; Bielen et al., 2007, Duffy et al., 2010, Kraus et al., 2015). All three Brachyury paralogs were detected in hypostome of hydrant with overlapping patterns (Figure 6d). As in *Hydra* (Bielen et al., 2007), overlapping expression domains of Brachyury paralogues in hypostomes of *D. pumila* hydranths suggest occurred subfunctionalization (Force et al., 1999).

In case of subfunctionalization, combined expression patterns of gene duplicates correspond approximately to that of the ancestral gene and reflects its original function (Force et al., 1999). We suggest that in a hydrozoan hydranth, the original function of Brachyury is associated with a specification of an oral domain (a hypostome), but not with a particular germ layer. Indeed, association between Brachyury orthologs and germ layers of hypostome is inconsistent among hydrozoans. For example, Brachyury1 ortholog is predominantly expressed in the ectoderm in *H. echinata* (Duffy et al., 2010) and in the endoderm in *Hydra* (Bielen et al., 2007), but in both germ layers in *D. pumila* (Figure 6d). On the contrary to *D. pumila* (Figure 6d), in *Hydra*, Brachyury2 ortholog is expressed in the ectoderm of the hypostome (Bielen et al., 2007). It indicates species-specific spatial regulation of Brachyury genes expression in hypostomes of hydrozoan hydranths.

Unfortunately, there are little studies of Brachyury expression in hydrozoan medusae and complex colonies. In case of medusae, it was shown that Brachyury3 is expressed in the developing bell and muscles of medusa buds and at the tip of the manubrium in the medusa of *P. carnea* (Spring et al., 2002). In *C. hemisphaerica*, ChBra1 is also expressed in medusa buds and at the tip of the manubrium (Kraus et al., 2015), and moreover according to transcryptome analysis, ChBra3 is also a medusa-specific gene (Leclère et al., 2019). Still, we need more data on Brachyury expression in hydrozoan medusae.

In the complex hydrozoan colonies, Brachyury expression was examined in the shoot growing tip of *D. pumila* (Bagaeva et al., 2019) (Figure 6c). While DpBra3 expression was not detected in shoot growing tip, DpBra1 and DpBra2 are strongly expressed in its apical ectoderm (Figure 6c). DpBra1 is expressed at the top of the developing shoot growth tip, but DpBra2 expression is absent from its central part and is associated with the formation of hydrant primordia at its opposite sides. Thus, DpBra2 seems to acquire a novel function in the hydranth primordia in *D. pumila*.

To further address the functional differences of Brachyury paralogs in *D. pumila*, we employed the *Xenopus* animal cap assay. Ability of *Brachyury* gene product of a wide range of species starting from the filasterean *Capsaspora owzcarzaki* to induce mesoderm when injected into the frog embryo (Marcellini et al., 2003; Sebe-Pedros et al., 2013) suggests functional conservation. It was shown that HyBra1 promotes the formation of mesoderm in the *Xenopus* animal cap whereas HyBra2 demonstrate a neural-inducing activity (Bielen et al., 2007).

In our study, DpBra1 induced expression of both early and late mesodermal marker genes, MyoD (myod.S) and muscle actin (actc1.L) (Figure 8a, c). These genes are downstream targets of *Xenopus* Brachyury (Sebe-Pedros et al., 2013), thus, DpBra1 roughly mimics the endogenous *Xenopus* Brachyury function. DpBra2 revealed neither mesoderm-inducing activity nor, in contrast to data obtained in Hydra (Bielen et al., 2007), a neural-inducing activity. As DpBra2, DpBra3 also didn’t show neural- or mesodermal-inducing activity (Figure 8a, c). These data indicate that functional divergence is characteristic for Brachyury2 and Brachyury3.

The specificity of Brachyury function is mostly defined by the N- and C-terminal domains, but not by the central T-box (Marcellini, 2006; Sebe-Pedros et al., 2013). In line with previous studies (Bielen et al., 2007) and our qRT-PCR data (Figure 8), hydrozoan Brachyury1 orthologs demonstrate high conservation of their overall protein sequence (Figure 2c), which is consistent with their high functional conservation. In Brachyury2 and Brachyury3, compared to T-boxes, N- and C-terminal domains show lesser amino acid identity to the ancestral gene and have lost ancestral C-terminal repression domain R1 (Figure 2a-c). These differences in terminal domains could be responsible for the neo- and subfunctionalization of Brachyury2 and Brachyury3 in hydrozoans, even though it was suggested that neo- and subfunctionalization are mainly due to mutations in regulatory sequences, rather than mutations in the coding sequence (Jayamaran et al., 2022).

Taken together, our data indicate two duplication events of Brachyury in the hydrozoan lineage. Brachyury1 is the most conservative duplicate, both on the functional and sequence levels. In studied hydrozoans and in *D. pumila* in particular, it is supposed to preserve its ancestral function as a crucial component of axis formation and patterning. Brachyury2 reveals features of both sub- and neofunctionalization: in studied hydrozoans, it seems to be a component of the axial patterning network together with Brachyury1 throughout the cnidarian life cycle. However, it also specificly marks forming hydranth primordia in a complex hydrozoan colony of *D. pumila*. Brachyury3 is the paralog that reveals strong divergence in sequence and functions among hydrozoans and reveals signs of both neo- and subfunctionalization in a species-specific manner. These findings extend the understanding of the function and evolution of Brachyury gene family in cnidarians and highlight the functional plasticity of Brachyury genes. Obtained data reaffirm that the border between sub- and neofunctionalization is often indistinct and the specific outcome of duplication is spatio- and temporal-dependent (He, Zhang, 2005). The case of subsequent gene duplication in hydrozoans seems to be a promising model for studies on post-duplication scenarios.

## Methods

### Animals and Sampling

Sampling of *D. pumila* colonies and experimental procedures over *D. pumila* embryos were performed at the Pertsov White Sea Biological Station (Lomonosov Moscow State University) (Kandalaksha Bay; 66°340 N, 33°080 E) during the period of *D. pumila* sexual reproduction (June-July). Sexually mature colonies were kept in natural seawater at +10-12°C. Whole-mount observations were made under a stereomicroscope Leica M165C.

### Chemical treatment

To activate/inhibit cWnt signaling, gastrulating embryos were treated with 0.5/1/2.5 μM 1-Azakenpaullone (Sigma, Canada/China) or 1/2.5/10 μM iCRT-14 (Sigma, USA/China) respectively. Stock solutions were prepared with DMSO at 10mM, aliquoted and stored at −20°C. Working solutions were prepared before use by dilution of stock solutions in filtered seawater (FSW) to the final concentration. Control embryos were exposed to 0.1% DMSO in FSW. Working solutions were refreshed daily. Incubation was performed in the dark.

### Data sources and Transcryptome assembly

To analyse phylogenetic relationships within the brachyury gene family, we surveyed 28 metazoan species. Gene sequences were obtained from several sources (Supplementary Table 1). Bilaterian, ctenophore, placozoan and anthozoan sequences were obtained from nucleotide collection of NCBI database. Some assembled cnidarian transcriptomes were downloaded from public databases at NCBI (*Aurelia aurita, Morbakka virulenta, Nemopilema nomurai, Podocoryna carnea, Lucernaria quadricornis, Tripedalia cystophora* (Khalturin et al., 2019), *Dynamena pumila* (Kupaeva et al., 2020), *Polypodium hydriforme* (Shpirer et al, 2014) or other web-sites (*Clytia hemisphaerica* (Leclère et al, 2019), *Hydractinia symbiolongicarpus* (https://research.nhgri.nih.gov/hydractinia/)). Transcriptomes of *Craspedacusta sowerbii* and *Margelopsis haeckelii* were newly assembled by ourself. Data for *Margelopsis haeckelii* were collected and sequenced *de novo* and are available in our lab. Read quality control was performed with fastp (v.0.20.0) software (Chen et al., 2018). *De novo* transcriptomes were assembled with rnaSPAdes (v.3.13.1) (Bankevich et al., 2012) software. Quality of assembly was assessed using BUSCO v.3.0.2 with metazoan database (Seppey et al., 2019).

### Phylogenetic analyses

Brachyury genes ABJ16449.1 and JAC85032.1 of *C. hemisphaerica* were used as queries for local tblastx search of Brachyury genes in *D. pumila* transcriptome. Using three obtained sequences of *D. pumila* Brachyury-like genes as queries, we searched for Brachyury-like genes in 10 other medusozoan transcriptomes. We surveyed 12 medusozoan transcriptomes in total (Supplementary Table 1). Obtained nucleotide sequences of medusozoan Brachyury-like genes together with sequences of bilaterian, ctenophore, placozoan and anthozoan Brachyury genes were translated with NCBI ORFfinder. Amino acid sequence alignments and phylogenetic analysis were performed with MUSCLE algorithm in MUSCLE software (v3.8.31) (Edgar et al., 2004). Sequences of Tbx genes were selected as an outgroup. Sequence alignments were trimmed by removing poorly aligned regions using TrimAL tool, v.1.2rev59 (Capella-Gutiérrez et al., 2009). A heuristic approach “automated1” was used to select the best automatic method to trim our alignments. Phylogenetic analysis was performed with Maximum Likelihood using IQTree v.2.0-rc2 software (Minh, et al., 2020). The JTT+R5 model was found to be optimal. To assess branch supports, bootstrap values were calculated running 1000 replicates using ultrafast bootstrap (UFBoot) (Hoang et al, 2018). Trees were visualized in FigTree v1.4.4 software. Obtained phylogenetic trees were processed with Adobe Illustrator CC. No corrections were made to the tree topology and the branch lengths.

To analyze functional domains of the hydrozoan Brachyury, selected protein sequences were scanned against Pfam hidden Markov model (HMM) database using *hmmscan* of HmmerWeb v.2.41.1 (Potter et al., 2018). Identification of the conserved R1 domain within the hydrozoan Brachyury was carried out using ClustalW sequence alignment service (Madeira et al., 2019) with the R1 domain in HyBra1 (Bielen et al., 2007) as a query. The domain architecture of proteins was visualized using Pfam (Mistry et al., 2020). Multiple sequence alignment and calculation of the identity matrix of hydrozoan Brachyury proteins and T-boxes of *D. pumila* Brachyury were conducted using ClustalW with default settings and shaded using BOXSHADE 3.21.

### *D. pumila* genes isolation, PCR, and antisense RNA probe synthesis

cDNA expression library was prepared by the SMART approach from total embryonic RNA with a Mint cDNA synthesis kit (Evrogen, Russia). cDNA gene fragments were isolated from the library by PCR with gene-specific primers (see Table 1). Primers were designed based upon sequences obtained from the sequenced transcriptome (Illumina) of D. pumila (Kupaeva et al., 2020). Amplified fragments were cloned into the pAL-TA vector (Evrogen, Russia). Digoxygenine□labeled antisense RNA probes were generated from gene fragments, which were amplified from plasmids with *D. pumila* genes.

**Table 1:**
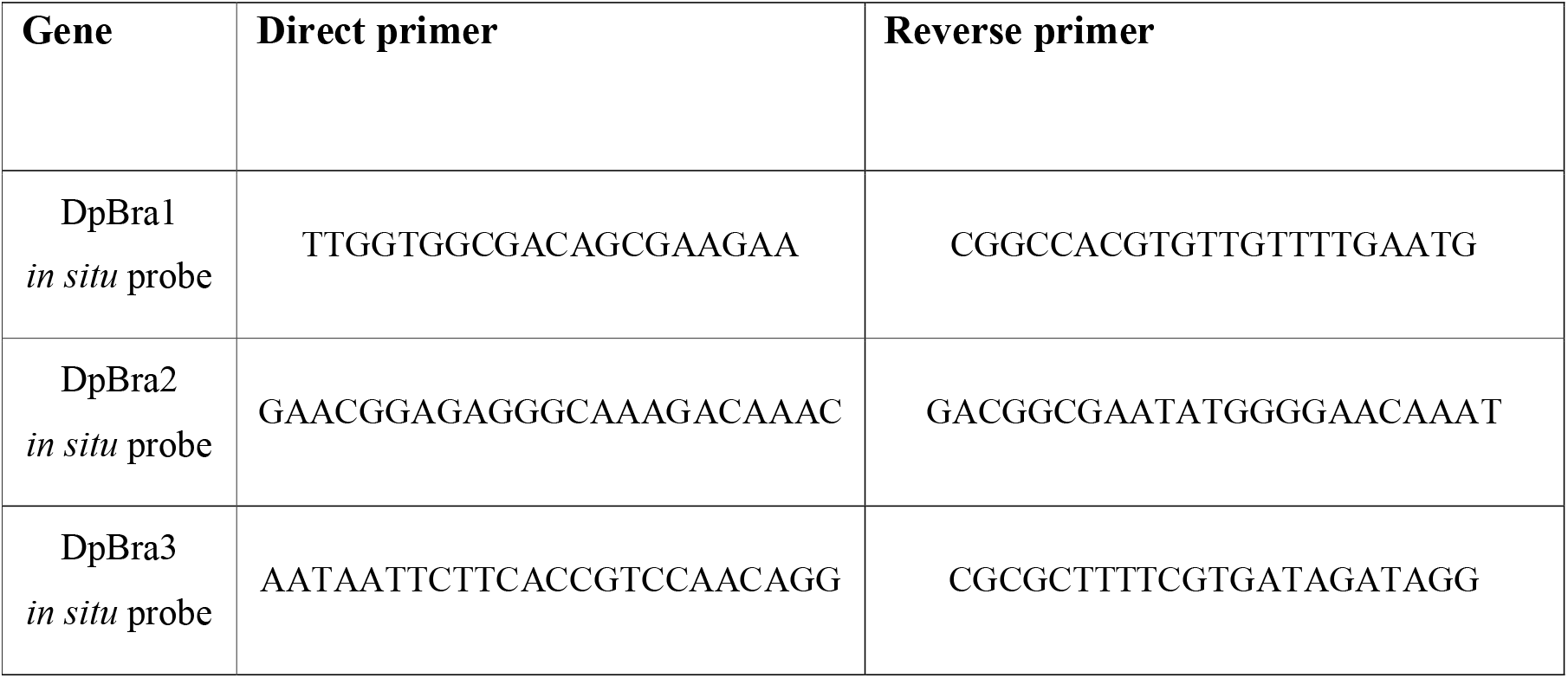
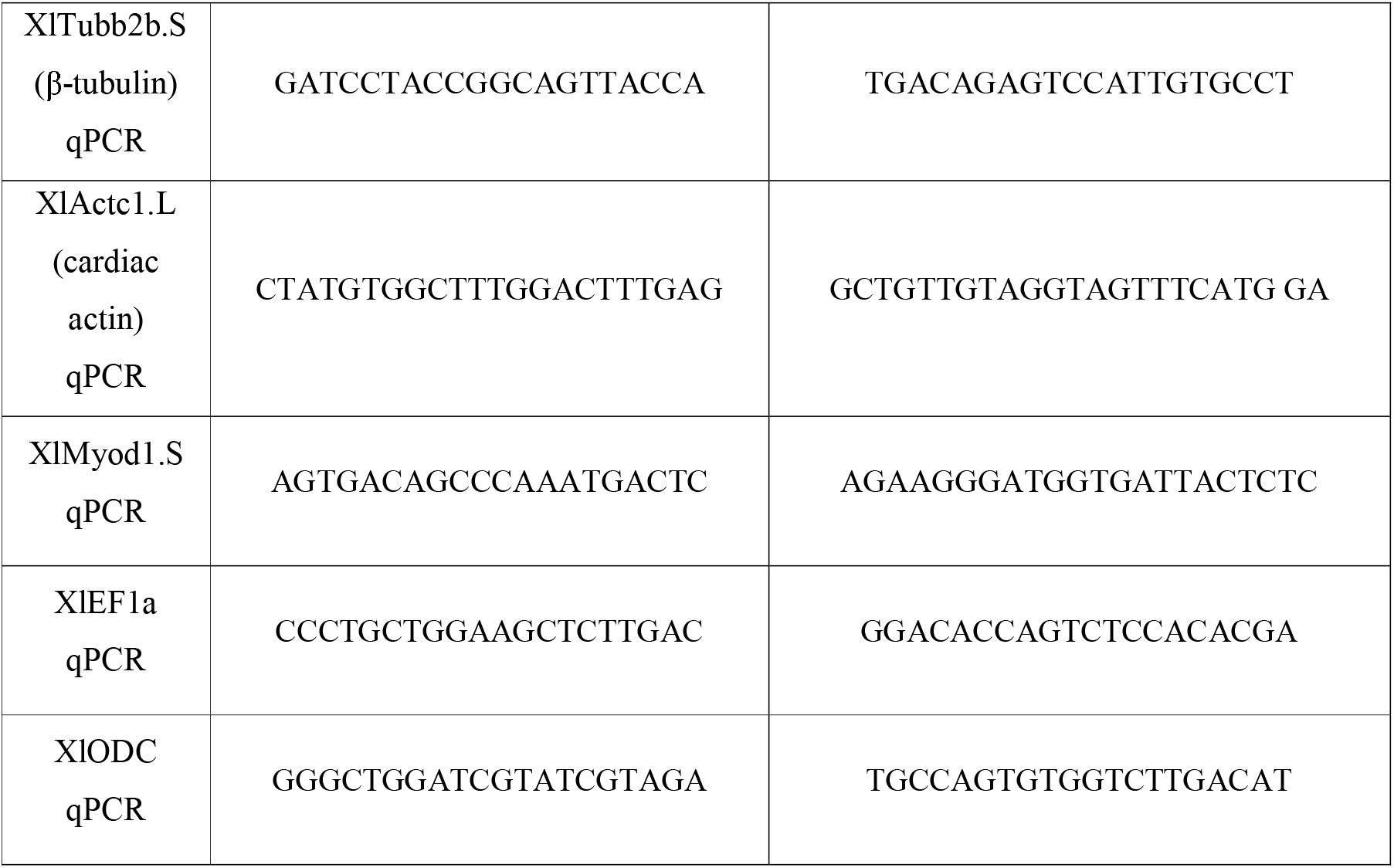
PCR and qPCR primers used in this study.

### *In situ* hybridization

The *in situ* hybridization protocol was performed as previously described in Bagaeva et al., 2019 for *D. pumila* shoots and hydranths and in Vetrova et al., 2021 for *D. pumila* embryos. An urea-based in situ hybridization method was used for the hydranths (Sinigaglia et al., 2017).

Shoots were fixed with 0.2% glutaraldehyde/4%formaldehyde in FSW for 1 minute and then for an additional hour with 4% formaldehyde in FSW. Samples were washed with PTw (1x PBS with 0.1% Tween 20) thrice and stored in 100% methanol no more than overnight at -20°C until hybridization. Embryos were fixed with 4% paraformaldehyde in FSW overnight at +4°C, rinsed with PBS, and stored at -20°C in 100% methanol until hybridization.

Samples were rehydrated with PTw and treated with proteinase K (80 μg/ml, 22°C) for 1-3 minutes. To inactivate the endogenous alkaline phosphatase and avoid a false positive result, samples were heated at +80°C for 30 minutes. Hybridization was performed at 62°C (shoots) or 58°C (embryos) with digoxigenin-labelled antisense RNA probes (1ng/μL). Anti[DIG alkaline phosphatase-conjugated antibody (Roche; 1/2000 diluted) and NBT/BCIP substrate (Roche) were used to detect the probe. Stained samples were washed with PTw and methanol to reduce background staining and mounted in glycerol (87%).

Several specimens were treated with Murray’s Clear solution (2:1 mixture of benzyl benzoate and benzyl alcohol) to achieve optical tissue clearing. Several specimens were embedded into Technovit resin. Sections (5-7 μm thick) were cut using Reichert-Jung (Leica) Ultra-cut 701701 ultramicrotome (Reichert-Jung, Austria). Imaging of samples was conducted using Leica M165C microscope (Leica, German) equipped with Leica DFC420C (5.0MP) digital camera.

### Animal cap assay

Wild-type *Xenopus laevis* were obtained from the European Xenopus Resource Centre (EXRC) at University of Portsmouth, School of Biological Sciences, UK, or Xenopus 1, USA. Frog maintenance and care was conducted according to standard procedures in the AquaCore facility, University Freiburg, Medical Center (RI_00544) and based on recommendations provided by the international Xenopus community resource centers NXR and EXRC as well as by Xenbase (http://www.xenbase.org/, RRID:SCR_003280). This work was done in compliance with German animal protection laws and was approved under Registrier-Nr. G-18/76 by the state of Baden-Württemberg.

*X. laevis* eggs were collected and in vitro-fertilized, then cultured and microinjected by standard procedures (Sive et al., 2010). Embryos were injected two times/embryo with mRNAs at two-cell or four-cell stage using a PicoSpritzer setup in 1/3x Modified Frog Ringer’s solution (MR) with 2.5% Ficoll PM 400 (GE Healthcare, #17-0300-50), and were transferred after injection into 1/3x MR containing Gentamycin. Drop size was calibrated to about 7–8nL per injection. Injected or uninjected (control) embryos were cultured until st. 8. Animal caps were dissected in 1x Modified Barth’s solution (MBS) and transferred to 0.5x MBS + Gentamycin. 10-15 organoids were collected in TRIzol per condition and experiment.

Full-length *D. pumila* Brachyury sequences were amplified from cDNA library and cloned into pCS2+8 plasmid. pCS2+8 was a gift from Amro Hamdoun (Addgene plasmid #34931; http://n2t.net/addgene:34931; RRID:Addgene_34931) (Gokirmak et al., 2012). mRNAs were prepared using the Ambion mMessage Machine kit using Sp6 (#AM1340) supplemented with RNAse Inhibitor (Promega #N251B) after plasmid linearization with Not1, and injected at 50ng/ul.

### RT-PCR

Total RNA was extracted using a standard Trizol (Invitrogen #15596026) protocol and used for cDNA synthesis with either iScript cDNA Synthesis Kit (Bio-Rad #1708891). qPCR-reactions were conducted using Sso Advanced Universal SYBR Green Supermix (Bio-Rad #172-5275) on a CFX Connect Real-Time System (Bio-Rad) in 96-well PCR plates (Brand #781366). Conventional PCR and gel-electrophoresis was conducted analogously on a S1000 Thermal cycler (Bior-Rad). See Table 1 for gene-specific primers. Expression values were normalized against two housekeeping control genes - EF1 and ODC (2^ΔΔCt^ method). Results are presented as means±□standard deviation (s. d.) of the relative fold change (rFC), which is a ratio of normalized mRNA level of the analysed gene expression in experimental group in comparison to control group.

### Statistical analysis

Statistical analysis of the normalized gene expression data after qRT-PCR was performed in GraphPad Prism 5 software. Normality of data distribution was checked by the Kolmogorov-Smirnov tests. Differences between groups were assessed with one-way ANOVA followed by Dunnet post hoc test. Significance is indicated by asterisks on the graphs. A P-value less than 0.05 was considered significant for all analysis. All experiments were designed with matched control conditions to enable statistical comparison. The n value is 7 for a control group. The n value for each experimental group is stated on graphs.

### Image processing

Pictures were edited with Adobe Photoshop CS6 programs. To achieve optimal exposure and contrast, alterations to the “Brightness’’, “Contrast”, “Exposure”, and “Levels” for the RGB channel were used. All tools were applied to the entire image, not locally.

## Data availability

Sequences obtained in this study have been deposited in GenBank under the following accession numbers: OP828770 – OP828776, OP902368, OP902367.

## Acknowledgements

We thank N.A. Pertsov White Sea Biological Station of Moscow State University for the help and support in obtaining samples and providing access to all required facilities and equipment of the “Center of Microscopy WSBS MSU”. We are grateful to Dr. Amro Hamdoun for pCS2+8 plasmid.

## Author contributions

A.A.V. conducted the experiments, performed a sequence analysis, performed a statistical analysis, interpreted the results, wrote the draft manuscript, and prepared the figures. D.M.K. assembled transcriptomes and performed a phylogenetic analysis. T.S.L. participated in the conduction of experiments. P.W. performed animal cap assay and quantitative RT-PCR. N.T. participated in data visualisation. S.V.K. conceptualized, designed and supervised the study, conducted the experiments, and interpreted the results. All authors read, revised and approved the final manuscript.

## Competing interests

The authors declare no competing interests.

**Supplementary Table 1.** List of the sequences analyzed in the study.

| Species | gene | Accession number | Source |
| --- | --- | --- | --- |
| *Acropora millepora* | Brachyury | KJ914894.1 | NCBI |
| *Aurelia aurita* | Brachyury 1 | GHAI01095397.1 | NCBI |
| *Aurelia aurita* | Brachyury 2 | GHAI01133198 | NCBI |
| *Aurelia aurita* | TBX4-5 | LN611631.1 | NCBI |
| *Beroe pileus* | Brachyury | CAE45766.1 | NCBI |
| *Branchiostoma floridae* | TBX1/10 | AAG34887.2 | NCBI |
| *Clytia hemisphaerica* | Brachyury 1 | ABJ16449.1 | NCBI |
| *Clytia hemisphaerica* | Brachyury 2 | JAC85032.1 | NCBI |
| *Clytia hemisphaerica* | Brachyury 3 | TCONS_00020933_XLOC_011741 | http://marimba.obs-vlfr.fr/node/237575 |
| *Clytia hemisphaerica* | TBX2-3 | LN611642.1 | NCBI |
| *Clytia hemisphaerica* | TBX4-5 | LN828922.1 | NCBI |
| *Craspedacusta sowerbii* | Brachyury 1 | OP902368 | NCBI |
| *Craspedacusta sowerbii* | Brachyury 3 | OP902367 | NCBI |
| *Danio rerio* | T-box transcription factor 16 | NP571133.1 | NCBI |
| *Danio rerio* | TBX6 | NP705952.2 | NCBI |
| *Drosophila melanogaster* | T-box protein H15 | CAA67304.1 | NCBI |
| *Dynamena pumila* | Brachyury 1 | MK005873.1 | NCBI |
| *Dynamena pumila* | Brachyury 2 | MK005874.1 | NCBI |
| *Dynamena pumila* | Brachyury 3 | OP828770 | NCBI |
| *Exaiptasia pallida* | Brachyury | XM021039556 | NCBI |
| *Homo sapiens* | TBX19 | NP005140.1 | NCBI |
| *Homo sapiens* | Brachyury (T) | CAA04938.1 | NCBI |
| *Homo sapiens* | TBX3 | AAC12947.1 | NCBI |
| *Homo sapiens* | TBX5 | AAC51644.1 | NCBI |
| *Homo sapiens* | TBX22 | NP001103348.1 | NCBI |
| *Homo sapiens* | TBX18 | CAB37937.1 | NCBI |
| *Hydra vulgaris* | Brachyury 1 | AY366371.1 | NCBI |
| *Hydra vulgaris* | Brachyury 2 | AY366372.1 | NCBI |
| *Hydractinia symbiolongicarpus* | Brachyury 1 | evgtrinLocDN7803c0g1t1 | https://research.nhgri.nih.gov/hydractinia/download/index.cgi?dl=s_ss |
| *Hydractinia symbiolongicarpus* | Brachyury 2 | evgtrinLocDN25577c0g1d264163t1 | https://research.nhgri.nih.gov/hydractinia/download/index.cgi?dl=s_ss |
| *Hydractinia symbiolongicarpus* | Brachyury 3 | evgtrinLocDN21629c0g1t2 | https://research.nhgri.nih.gov/hydractinia/download/index.cgi?dl=s_ss |
| *Lucernaria quadricornis* | Brachyury 2 | HAHD01071616 | NCBI |
| *Margelopsis haeckelii* | Brachyury 1 | OP828774 | NCBI |
| *Margelopsis haeckelii* | Brachyury 2 | OP828775 | NCBI |
| *Margelopsis haeckelii* | Brachyury 3 | OP828776 | NCBI |
| *Morbakka virulenta* | Brachyury 1 | GHAF01048756.1 | NCBI |
| *Morbakka virulenta* | Brachyury 2 | GHAF01045583.1 | NCBI |
| *Mus musculus* | Tbx2 | NP033350.2 | NCBI |
| *Mus musculus* | Tbx4 | NP035666.2 | NCBI |
| *Mus musculus* | Tbx6 | AAC53110.1 | NCBI |
| *Mus musculus* | Tbx10 | NP001001320.1 | NCBI |
| *Mus musculus* | Tbx1 | P70323.2 | NCBI |
| *Mus musculus* | Tbx20 | NP065242.1 | NCBI |
| *Mus musculus* | Tbx15 | O70306.2 | NCBI |
| *Nematostella vectensis* | Brachyury | AF540387.2 | NCBI |
| *Nematostella vectensis* | Tbx1 | AAQ23383.1 | NCBI |
| *Nemopilema nomurai* | Brachyury 1 | GHAR01071730.1 | NCBI |
| *Nemopilema nomurai* | Brachyury 2 | GHAR01060992 | NCBI |
| *Parasteatoda tepidariorum* | T-box protein H15-2 | BAD16721.1 | NCBI |
| *Podocoryna carnea* | Tbx4/5 | CAE45765.1 | NCBI |
| *Podocoryna carnea* | Brachyury 1 | GCHV01005394.1 | NCBI |
| *Podocoryna carnea* | Brachyury 2 | GCHV01013749.1 | NCBI |
| *Podocoryne carnea* | Brachyury (Brachyury 3) | AJ428494.1 | NCBI |
| *Polypodium hydriforme* | Brachyury | GBGH01023047.1 | NCBI |
| *Saccoglossus kowalevskii* | Tbx2/3 | NP001158392.1 | NCBI |
| *Stylophora pistillata* | Brachyury | XM022931758.1 | NCBI |
| *Tribolium castaneum* | Tbx20 | XP972670.3 | NCBI |
| *Trichoplax adhaerens* | Brachyury | CAD70269.1 | NCBI |
| *Trichoplax adhaerens* | Tbx2/3 | CAD70270.1 | NCBI |
| *Tripedalia cystophora* | Brachyury 2 | GHAQ01080409.1 | NCBI |
| *Tripedalia cystophora* | Brachyury 2 | GHAQ01019284.1 | NCBI |
| *Xenopus laevis* | VegT | AAB93301.1 | NCBI |

## References

1. Sebé-Pedrós, A. et al. The dynamic regulatory genome of Capsaspora and the origin of animal multicellularity. Cell. 165, 1224–1237 (2016).

2. Dobrovolskaia-Zavadskaia, N. Sur la mortification spontanee de la queuw chez la spuris nouveau et sur l’existence d’un caractere (facteur) hereditaire non viable. Compr. Soc. Biol. 97, 114–119 (1927).

3. Smith, J. Brachyury and the T-box genes. Curr. Opin. Genet. Dev. 7, 474-480 (1997)

4. Gluecksohn-Schoenheimer, S. The development of normal and homozygous brachy (T/T) mouse embryos in the extraembryonic coelom of the chick. Proc. Natl. Acad. Sci. USA. 30, 134–140 (1944).

5. Gruneberg, H. Genetical studies on the skeleton of the mouse. XXIII. The development of Brachyury and Anury. J. Embryol. Exp. Morphol. 6, 424–443 (1958).

6. Harada, Y., Yasuo, H., Satoh, N. A sea urchin homologue of the chordate Brachyury (T) gene is expressed in the secondary mesenchyme founder cells. Development 121, 2747–2754 (1995).

7. Technau, U. Brachyury, the blastopore and the evolution of the mesoderm. Bioessays 23, 788–794 (2001).

8. Yamada, A., Martindale, M. Q., Fukui, A., Tochinai, S. Highly conserved functions of the Brachyury gene on morphogenetic movements: insight from the early-diverging phylum Ctenophora. Dev. Biol. 339, 212–222 (2010).

9. Sebé-Pedrós, A. et al. Early evolution of the T-box transcription factor family. Proc Natl Acad Sci U S A. 110, 16050–16055 (2013).

10. Satoh, N., Tagawa, K., Takahashi, H. How was the notochord born?. Evol. Dev. 14, 56–75 (2012).

11. Bruce, A. E. E., Winklbauer, R. Brachyury in the gastrula of basal vertebrates. Mech. Dev. 163, 103625; 10.1016/j.mod.2020.103625 (2020).

12. Yasuoka, Y., Shinzato, C., Satoh, N. The Mesoderm-Forming Gene brachyury Regulates Ectoderm-Endoderm Demarcation in the Coral Acropora digitifera. Curr. Biol. 26, 2885–2892 (2016).

13. Lapébie, P. et al. Differential responses to Wnt and PCP disruption predict expression and developmental function of conserved and novel genes in a cnidarian. PLoS Genet. 10, e1004590; 10.1371/journal.pgen.1004590 (2014).

14. Schwaiger, M., Andrikou, C., Dnyansagar, R. et al. An ancestral Wnt–Brachyury feedback loop in axial patterning and recruitment of mesoderm-determining target genes. Nat Ecol Evol 6, 1921–1939 (2022).

15. Kiecker, C., Niehrs, C. A morphogen gradient of Wnt/beta-catenin signalling regulates anteroposterior neural patterning in Xenopus. Development 128, 4189–4201 (2001).

16. Darras, S., Gerhart, J., Terasaki, M., Kirschner, M., Lowe, C. J. β-catenin specifies the endomesoderm and defines the posterior organizer of the hemichordate Saccoglossus kowalevskii. Development 138, 959–970 (2011).

17. Peter, I. S., Davidson, E. H. A gene regulatory network controlling the embryonic specification of endoderm. Nature 474, 635–639 (2011).

18. Bagaeva, T. et al. β-catenin dependent axial patterning in Cnidaria and Bilateria uses similar regulatory logic. Preprint at https://www.biorxiv.org/content/10.1101/2020.09.08.287821v1 (2020).

19. Kraus, Y., Chevalier, S., Houliston, E. Cell shape changes during larval body plan development in Clytia hemisphaerica. Dev. Biol. 468, 59–79 (2020).

20. Scholz, C. B., Technau, U. The ancestral role of Brachyury: expression of NemBra1 in the basal cnidarian Nematostella vectensis (Anthozoa). Dev. Genes Evol. 212, 563–570 (2003).

21. Croce, J., Lhomond, G., Gache, C. Expression pattern of Brachyury in the embryo of the sea urchin Paracentrotus lividus. Dev. Genes Evol. 211, 617–619 (2001).

22. Shoguchi, E., Satoh, N., Maruyama, Y. K. A starfish homolog of mouse T-brain-1 is expressed in the archenteron of Asterina pectinifera embryos: possible involvement of two T-box genes in starfish gastrulation. Dev. Growth Differ. 42, 61–68 (2000).

23. Yuan, L., Wang, Y., Li, G. Differential expression pattern of two Brachyury genes in amphioxus embryos. Gene Expr. Patterns. 38, 119152; 10.1016/j.gep.2020.119152 (2020).

24. Yasuo, H., Satoh, N. An Ascidian Homolog of the Mouse Brachyury (T) Gene is Expressed Exclusively in Notochord Cells at the Fate Restricted Stage: (Ascidians/T (Brachyury) gene/sequence conservation/notochord cells/transient expression). Dev. Growth Differ. 36, 9–18 (1994).

25. Latinkić, B. V. et al. The Xenopus Brachyury promoter is activated by FGF and low concentrations of activin and suppressed by high concentrations of activin and by paired-type homeodomain proteins. Genes Dev. 11, 3265–3276 (1997).

26. Hayata, T., Kuroda, H., Eisaki, A., Asashima, M. Expression of Xenopus T-box transcription factor, tbx2 in Xenopus embryo. Dev Genes Evol. 209, 625–628 (1999).

27. Session, A. M. et al. Genome evolution in the allotetraploid frog Xenopus laevis. Nature. 538, 336–343 (2016).

28. Martin, B. L., Kimelman, D. Regulation of canonical Wnt signaling by Brachyury is essential for posterior mesoderm formation. Dev Cell. 15, 121–133 (2008).

29. Terazawa K., Satoh N. Spatial expression of the amphioxus homologue of Brachyury (T) gene during early embryogenesis of Branchiostoma belcheri. Dev. Growth Differ. 37, 395–401 (1995).

30. Terazawa K., Satoh N. Formation of the chordamesoderm in the amphioxus embryo: Analysis with Brachyury and fork head/HNF-3 genes. Dev Genes Evol. 207, 1–11 (1997).

31. Inoue, J., Yasuoka, Y., Takahashi, H., Satoh, N. The chordate ancestor possessed a single copy of the Brachyury gene for notochord acquisition. Zoological Lett. 3, 1–7 (2017).

32. Bielen, H. et al. Divergent functions of two ancient Hydra Brachyury paralogues suggest specific roles for their C-terminal domains in tissue fate induction. Development 134, 4187–4197 (2007).

33. Vonica, A., Gumbiner, B. M. Zygotic Wnt activity is required for Brachyury expression in the early Xenopus laevis embryo. Dev. Biol. 250, 112–127 (2002).

34. Momose, T., Houliston, E. Two oppositely localised frizzled RNAs as axis determinants in a cnidarian embryo. PLoS Biol. 5, e70; 10.1371/journal.pbio.0050070 (2007).

35. Momose, T., Derelle, R., Houliston, E. A maternally localised Wnt ligand required for axial patterning in the cnidarian Clytia hemisphaerica. Development. 135, 2105–2113 (2008).

36. Kunick, C., Lauenroth, K., Leost, M., Meijer, L., Lemcke, T. 1-Azakenpaullone is a selective inhibitor of glycogen synthase kinase-3 beta. Bioorg. Med. Chem. Lett. 14, 413–416 (2004).

37. Stukenbrock, H. et al. 9-cyano-1-azapaullone (cazpaullone), a glycogen synthase kinase-3 (GSK-3) inhibitor activating pancreatic beta cell protection and replication. J. Med. Chem. 51, 2196–2207 (2008).

38. Gonsalves, F. C. et al. An RNAi-based chemical genetic screen identifies three small-molecule inhibitors of the Wnt/wingless signaling pathway. Proc. Natl. Acad. Sci. U. S. A. 108, 5954–5963 (2011).

39. Vetrova, A. A. et al. From apolar gastrula to polarized larva: Embryonic development of a marine hydroid, Dynamena pumila. Dev Dyn. 251, 795–825 (2022).

40. Green, J. The animal cap assay. Methods Mol Biol. 127, 1–13 (1999).

41. Smith, J. C., Price, B. M., Green, J. B., Weigel, D., Herrmann, B. G. Expression of a Xenopus homolog of Brachyury (T) is an immediate-early response to mesoderm induction. Cell. 67, 79–87 (1991).

42. Gu, X. Evolution of duplicate genes versus genetic robustness against null mutations. Trends Genet. 19, 354–356 (2003).

43. Wagner, A. Gene duplications, robustness and evolutionary innovations. Bioessays 30, 367–373 (2008).

44. Perry, B. R., Assis R. CDROM: Classification of Duplicate gene RetentiOn Mechanisms. BMC Evol Biol. 16, 82 (2016).

45. Force, A et al. Preservation of duplicate genes by complementary, degenerative mutations. Genetics 151, 1531–1545 (1999).

46. Lynch, M., Conery, J. S. The evolutionary fate and consequences of duplicate genes. Science 290, 1151–1155 (2000).

47. Ohno, S. Evolution by Gene Duplication. (Springer-Verlag, 1970).

48. Krakauer, D. C., Nowak, M. A. Evolutionary preservation of redundant duplicated genes. Semin. Cell Dev. Biol. 10, 555–559 (1999).

49. Conant, G. C., Wolfe, K. H. Turning a hobby into a job: how duplicated genes find new functions. Nat Rev Genet. 9, 938–950 (2008).

50. Assis, R., Bachtrog, D. Neofunctionalization of young duplicate genes in Drosophila. Proc. Natl. Acad. Sci. U. S. A. 110, 17409–17414 (2013).

51. Gu, X., Zhang, Z., Huang, W. Rapid evolution of expression and regulatory divergences after yeast gene duplication. Proc. Natl. Acad. Sci. U.S.A. 102, 707–712 (2005).

52. Carroll, S. B. Chance and necessity: the evolution of morphological complexity and diversity. Nature 409, 1102–1109 (2001).

53. Larroux, C. et al. Genesis and expansion of metazoan transcription factor gene classes. Mol. Biol. Evol. 25, 980–996 (2008).

54. Schulte-Merker, S., Van Eeden, F. J., Halpern, M. E., Kimmel, C. B., Nusslein-Volhard, C. no tail (ntl) is the zebrafish homologue of the mouse T (Brachyury) gene. Development 120, 1009–1015 (1994).

55. Strong, C. F., Barnett, M. W., Hartman, D., Jones, E. A., Stott, D. Xbra3 induces mesoderm and neural tissue in Xenopus laevis. Dev Biol. 222, 405–419 (2000).

56. Kraus, Y.A., Markov, A.V. Gastrulation in Cnidaria: The key to an understanding of phylogeny or the chaos of secondary modifications?. Biol Bull Rev 7, 7–25 (2017).

57. Spring, J. et al. Conservation of Brachyury, Mef2, and Snail in the myogenic lineage of jellyfish: a connection to the mesoderm of bilateria. Dev Biol. 244, 372–384 (2002).

58. Servetnick, M. D. et al. Cas9-mediated excision of Nematostella brachyury disrupts endoderm development, pharynx formation and oral-aboral patterning. Development. 144, 2951–2960 (2017).

59. Tu, Q., Brown, C. T., Davidson, E. H., Oliveri, P. Sea urchin Forkhead gene family: phylogeny and embryonic expression. Dev Biol. 300, 49–62 (2006).

60. Grimmelikhuijzen, C. J., Westfall, J. A. The nervous systems of cnidarians. EXS. 72, 7–24 (1995).

61. Arnold, S.J. et al. Brachyury is a target gene of the Wnt/beta-catenin signaling pathway. Mech Dev. 91, 249–258 (2000).

62. Duffy, D. J., Plickert, G., Kuenzel, T., Tilmann, W., Frank, U. Wnt signaling promotes oral but suppresses aboral structures in Hydractinia metamorphosis and regeneration. Development. 137, 3057–3066 (2010).

63. Kraus, J. E., Fredman, D., Wang, W., Khalturin, K., Technau, U. Adoption of conserved developmental genes in development and origin of the medusa body plan. Evodevo. 6, 23 (2015).

64. Leclère, L. et al. The genome of the jellyfish Clytia hemisphaerica and the evolution of the cnidarian life-cycle. Nat. Ecol. Evol. 3, 801–810 (2019).

65. Bagaeva, T. S. et al. сWnt signaling modulation results in a change of the colony architecture in a hydrozoan. Dev. Biol. 456, 145–153 (2019).

66. Marcellini, S., Technau, U., Smith, J. C., Lemaire, P. Evolution of Brachyury proteins: identification of a novel regulatory domain conserved within Bilateria. Dev Biol. 260, 352–361 (2003).

67. He, X., Zhang, J. Rapid subfunctionalization accompanied by prolonged and substantial neofunctionalization in duplicate gene evolution. Genetics 169, 1157–1164 (2005).

68. Marcellini, S. When Brachyury meets Smad1: the evolution of bilateral symmetry during gastrulation. Bioessays. 28, 413–420 (2006).

69. Jayaraman, V., Toledo-Patiño, S., Noda-García, L., Laurino, P. Mechanisms of protein evolution. Protein Sci. 31, e4362; 10.1002/pro.4362 (2022).

70. Khalturin, K. et al. Medusozoan genomes inform the evolution of the jellyfish body plan. Nat. Ecol. Evol. 3, 811–822 (2019).

71. Kupaeva, D., Konorov, E., Kremnyov, S. De novo transcriptome sequencing of the thecate colonial hydrozoan, Dynamena pumila. Mar. Genomics. 51, 100726; 10.1016/j.margen.2019.100726 (2020).

72. Shpirer, E. et al. Diversity and evolution of myxozoan minicollagens and nematogalectins. BMC Evol. Biol. 14, 205 (2014).

73. Hydractinia Genome Project Portal https://research.nhgri.nih.gov/hydractinia/ (2022)

74. Chen, S., Zhou, Y., Chen, Y., Gu, J. fastp: an ultra-fast all-in-one FASTQ preprocessor. Bioinformatics. 34, i884–i890 (2018).

75. Bankevich, A. et al. SPAdes: a new genome assembly algorithm and its applications to single-cell sequencing. J. Comput. Biol.19, 455–477 (2012).

76. Seppey, M., Manni, M., Zdobnov, E. M. BUSCO: Assessing Genome Assembly and Annotation Completeness in Gene Prediction. Methods in Molecular Biology. Volume 1962 (ed. Kollmar, M.) 227–245 (Humana, 2019).

77. Edgar, R. C. MUSCLE: multiple sequence alignment with high accuracy and high throughput. Nucleic Acids. Res. 32, 1792–1797 (2004).

78. Capella-Gutiérrez, S., Silla-Martínez, J. M., Gabaldón, T. trimAl: a tool for automated alignment trimming in large-scale phylogenetic analyses. Bioinformatics. 25, 1972[1973 (2009).

79. Minh, B. Q. et al. IQ-TREE 2: New Models and Efficient Methods for Phylogenetic Inference in the Genomic Era. Mol Biol Evol. 37, 1530–1534 (2020).

80. Hoang, D. T., Chernomor, O., von Haeseler, A., Minh, B. Q., Vinh, L. S. UFBoot2: Improving the Ultrafast Bootstrap Approximation. Mol. Biol. Evol. 35, 518–522 (2018).

81. Potter, S. C. et al. HMMER web server: 2018 update. Nucleic Acids Res. 46, W200–W204 (2018).

82. Madeira, F. et al. The EMBL-EBI search and sequence analysis tools APIs in 2019. Nucleic Acids Res. 47, W636–W641 (2019).

83. Mistry, J. et al. Pfam: The protein families database in 2021. Nucleic Acids Res. 49, D412–D419 (2021).

84. Nieuwkoop, P.D., Faber, J. Normal Table of Xenopus Laevis (Daudin) A Systematical & Chronological Survey of the Development from the Fertilized Egg till the End of Metamorphosis (Garland Pub., 1994).

85. H. L. Sive, R. M. Grainger, R. M. Harland, Microinjection of xenopus oocytes. Cold Spring Harb. Protoc. 5 (2010), doi:10.1101/pdb.ip81.

86. Gökirmak, T. et al. Localization and substrate selectivity of sea urchin multidrug (MDR) efflux transporters. J Biol Chem. 287, 43876–43883 (2012).

